# An oncoembryology approach uncovers SoxC-driven regulation of colon development and cancer

**DOI:** 10.1101/2024.10.08.617191

**Authors:** Tosca Dalessi, Tomas Valenta, Giulia Moro, Michael David Brügger, Ab Meijs, Barbara M. Szczerba, Tamara Brunner, Simon Lampart, Marc Nater, Randall J. Platt, Michael Scharl, Maries van den Broek, Isabelle C. Arnold, Hassan Fazilaty

## Abstract

Reactivated embryonic programs are associated with cancer progression, yet their role and regulatory mechanisms remain poorly understood. In this study, we introduce ‘oncoembryology,’ an approach that systematically compares embryonic and cancerous tissues to identify shared molecular programs and assess their functional relevance in disease. Applying this strategy to colorectal cancer, we identified SoxC transcription factors (Sox4, Sox11, Sox12) as critical regulators of both embryonic development and tumorigenesis. SoxC transcription factors regulate diverse downstream targets, including Tead2, Mdk, and Klf4, thereby regulating crucial steps of colon development. Abrogating SoxC function in murine models reduced tumor growth and prevented liver metastasis. Concordantly, a SoxC-driven oncoembryonic gene signature correlated with poor survival in colorectal cancer patients, underscoring the therapeutic potential of targeting SoxC-regulated pathways in cancer treatment.

## Introduction

The phenotypical parallels between cancer cells and embryonic cells, first observed by pathologists such as Virchow, Lobstein, and Recamier in the 19th century, have recently been substantiated at the molecular level, revealing shared transcriptional signatures and genetic programs (Virchow, 1859; Krebs, 1947; Gold and Freedman, 1965; Nieto et al., 1994; Weinberg, 1996; Hanahan and Weinberg, 2011; Nieto, 2013; Aiello and Stanger, 2016; Hanahan, 2022; Fazilaty and Basler, 2023). While the controlled reactivation of embryonic programs is crucial for successful tissue regeneration, improper regulation or failure to silence these programs can lead to pathologies including malignant cancers (Fazilaty and Basler, 2023). Although the reactivation of *oncoembryonic* programs is a recognized phenomenon, the underlying mechanisms and their functional consequences remain poorly understood. Previous research, beyond a few well-known processes such as *epithelial-mesenchymal plasticity*, has primarily correlated embryonic genes expression and cancer progression without conducting simultaneous functional analyses in the embryo to clarify the actual roles these programs play in both contexts (Fazilaty and Basler, 2023; Mustata et al., 2013; Baulies et al., 2024; AR Moorman et al., 2023; Burdziak et al., 2023; Fazilaty et al., 2019; Youssef et al., 2024; Cano et al., 2000; Ocaña et al., 2012; Thiery et al., 2009). This gap in understanding underscores the need for research to unravel the role of oncoembryonic programs in cancer.

In this study, we adopted a systematic approach, first identifying common genetic components between embryonic development and cancer of the colon, then conducting functional analysis within the controlled embryonic context to uncover key regulatory mechanisms. The embryonic environment offers a robust platform for isolating functional components that are difficult to dissect in the complex tumor setting. Once validated, we returned to the cancer model to assess their relevance to disease progression and therapeutic potential. We term this reiterative approach ‘oncoembryology.’ Through this, we identified SoxC transcription factors (TFs) as key regulators of both embryonic colon development and colon cancer progression. By studying SoxC TFs in their native embryonic context, we pinpointed essential downstream factors as potential therapeutic targets in colon cancer. This work highlights critical aspects of oncoembryonic programs and demonstrates a proof-of-concept for strategies that could translate into more effective cancer treatments.

## Results

### SoxC transcription factors are oncoembryonic regulators involved in colorectal cancer

Colorectal cancer (CRC) is the top cause of cancer death in young adults, and its prevalence is increasing. Currently, patients with metastatic cancer have poor survival rates, making it clear that new therapies are urgently needed (Biller and Schrag, 2021; Siegel et al., 2024; Vuik et al., 2019). Leveraging the oncoembryology concept, we systematically compared the cell and molecular profiles of the colon during development (embryonic hindgut), healthy adult colon, and CRC. To faithfully model the conventional metastatic disease in mice, we employed colonoscopy-guided submucosal injection of organoids in syngeneic mice (Borrelli et al., 2024; Roper et al., 2018), using organoids harboring mutations in *Apc, Tp53, Kras*, and *Smad4* to enhance tumorigenesis and invasiveness, reflecting the common mutational landscape of CRC in patients (Fang et al., 2021; Hashimoto et al., 2024; Nakayama and Oshima, 2019; Phipps et al., 2013) (Figures S1A-F).

As a first step, we sought to identify the genes specifically expressed both in the embryonic hindgut and in CRC - the oncoembryonic genes. To this end we compared single-cell RNA sequencing (scRNAseq) data from the mouse developing hindgut at embryonic day (E) 14.5, healthy adult colon, and advanced colon tumors. We found two epithelial cell clusters enriched in tumors (Figure 1A). The first cluster corresponded to cells with embryonic features and contained most embryonic cells (cluster 1, oncoembryonic). Validating the classification of these as oncoembryonic cells, the cells expressed previously reported oncoembryonic markers such as *Clu* and *Tacstd2* (Figures S2A-B) (Bala et al., 2023; Fazilaty et al., 2021; Fazilaty and Basler, 2023; Mustata et al., 2013). Functional gene ontology analysis of the highly expressed genes in oncoembryonic cells pointed to processes like “integrin cell surface interactions”, “DNA damage-induced apoptosis regulation”, “macrophage chemotaxis” and “nucleosomal DNA binding” (Figure S2C). Functional protein association network analysis (Szklarczyk et al., 2023) of the proteins encoded by the highly expressed genes in oncoembryonic cells showed two main branches linked via proteins such as Spp1, Mdk and Fn1 (Figure S2D), suggesting their importance as drivers of tumor progression. The second tumor-specific cluster expressed tuft cell markers such as Pou2f3 and Dclk1 (cluster 2; tuft-like tumor, Figure 1A, and Figures S2A-B). However, this cluster did not contain any embryonic cell. Genes expressed in tuft-like tumor cells were linked to immune regulation and cellular signaling, in line with the known function of tuft cells (Figure S2E).

**Figure 1.**
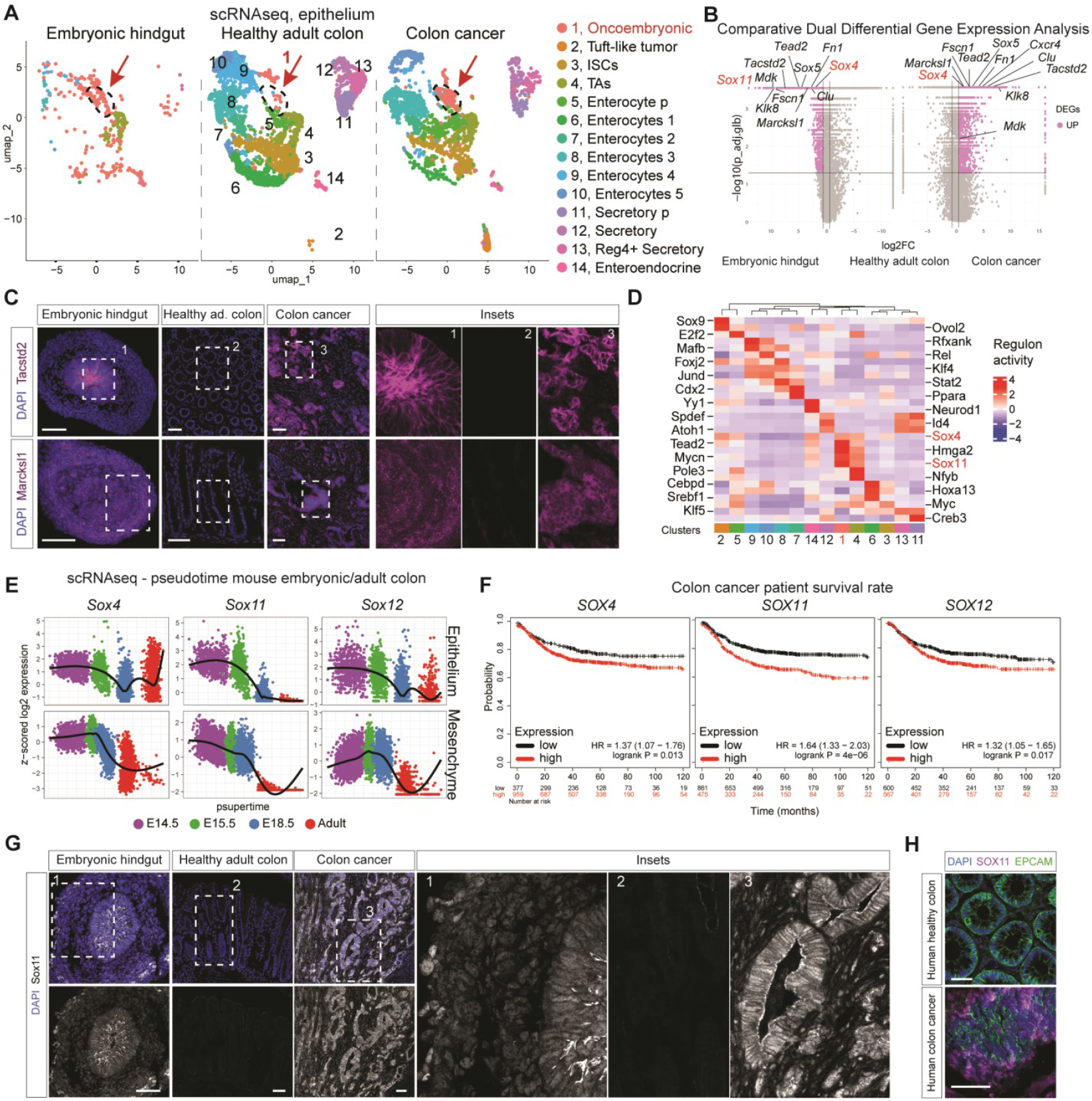
Reactivation of SoxC embryonic transcription factors in CRC is correlated with poor patient survival. (A) UMAP plot of integrated single-cell RNA sequencing (scRNAseq) datasets of mouse embryonic developing colon (hindgut), healthy adult colon and colon cancer (derived from AKPS organoid-based model). (B) Volcano plots of dual differential gene expression analysis comparing epithelial cell gene expression from scRNAseq data of embryonic hindgut and colon cancer to the healthy adult colon. Highlighted genes are examples of common genes in the embryonic and cancer cells. (log2FC (log2 fold change) > 0.6, p_adj.glb (p value adjusted global) < 0.05). Differential expression and p values were calculated via hierarchical permutation tests. (C) Immunofluorescent (IF) staining on embryonic hindgut, healthy adult colon and colon cancer for Tactsd2 and Marcksl1 (magenta). Blue DAPI. Scale bar: 50 μm. (D) Regulatory network inference using SCENIC analysis on the integrated epithelial scRNAseq data (from 1A). (E) Expression values of SoxC genes are shown as psupertime value learned for each cell (X-axis) and Z-scored log2 gene expression values (Y-axis) across three embryonic and one adult time-points. (F) Kaplan Meier plots showing the inverse correlation between SOXC family gene expression and CRC patients’ relapse-free survival. Hazard ratio (HR) and logarithmic ranked p Value (logrank P) were analyzed to infer the significance of the differences. Numbers below each graph represent number of patients at risk in any given time (months), black for low expression and red for high expression of each gene. The cut-off is automatically calculated based on the best performing threshold. (G) IF staining on embryonic hindgut, healthy adult colon and colon cancer for Sox11 (grey). Blue DAPI. Scale bar: 50 μm. (H) IF staining on human healthy and malignant colon for EPCAM (epithelial marker, green) and SOX11 protein (magenta). Blue DAPI. Scale bar: 50 μm.

To explore the full extent of embryonic-cancer overlap, we performed a comparative dual differential expression analysis, comparing the transcriptomes of all epithelial cells from embryonic hindgut and colon cancer cells to that of healthy adult colon epithelial cells. We identified over 1,500 oncoembryonic genes that were expressed in both embryonic and cancerous states, with minimal or no expression in healthy adult tissue (Figure 1B), mainly involved in processes such as “DNA replication”, “chromatin remodeling” “RNA processing” and “transcription regulation” (Figures S3A-B). Notably, genes like *Clu, Tacstd2, Marcksl1, Tead2* and *Sox4* were prominently expressed in oncoembryonic cells (Figures 1B-C). Regulatory network inference predicted strong activity of regulons driven by TFs including Sox4, Sox11, Mycn, Tead2 and Hmga2 in oncoembryonic cells (Figures 1D and S3C). Interestingly, Sox4 and Sox11, which together with Sox12 form the SoxC subfamily, are specifically expressed in embryonic epithelium and mesenchyme (Fig 1E).

To investigate the impact of SoxC gene reactivation in tumors, we assessed the relationship between elevated expression of these factors and CRC patient survival. Strikingly, high expression of SoxC TFs in human CRC tissues correlated with poor patient survival (Figure 1F). Compared to healthy adult tissues, where only low Sox4 levels are detectable, strong expression of both Sox4 and Sox11 is reactivated in tumors (Figures 1G-H and S3D-F). Taken together, these data led us to further investigate the role of SoxC TFs as master regulators of an oncoembryonic program in colorectal cancer.

### Studying SoxC in the embryo reveals its *bona fide* genetic program

The embryonic tissue provides a more stable and reproducible system for studying SoxC function than the complex tumor environment. Therefore, to discern the *bona fide* genetic program orchestrated by the SoxC subfamily, we first sought to understand the functional role of these TFs in embryonic development.

Using a conditional mouse model, we induced simultaneous ubiquitous knockout of all three SoxC TFs (SoxC-KO) to avoid complications due to redundancy among family members (Bhattaram et al., 2010; Miao et al., 2019). Recombination was induced at E10.5, when SoxC expression peaks in the developing intestine (Figure S4A). SoxC-KO embryos displayed significant developmental abnormalities such as underdeveloped limbs, malformed eyes as previously described (Bhattaram et al., 2010). In addition, we observed shorter intestinal length in KO embryos (Figures 2A and S4B-C). In the intestine, Dalessi et al., 2024 (preprint) SoxC-KO embryos exhibited decreased stromal width and fewer mesenchymal cells between the epithelium and the smooth muscle layer (Figures 2B-C and S4D). Taken together these results confirm the pivotal role of SoxC in development.

**Figure 2.**
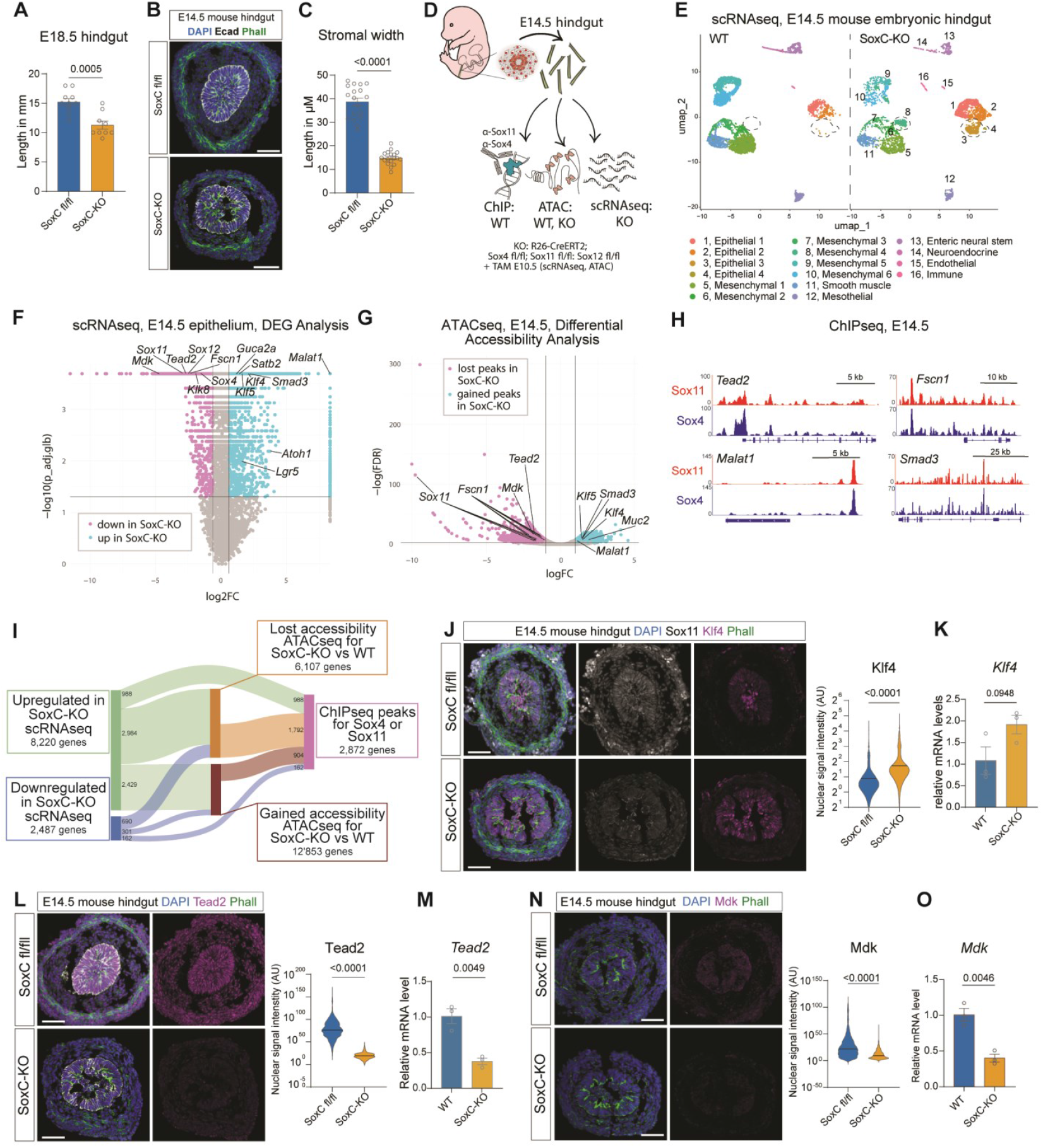
Knock-out model and multi-omics approach to identify SoxC-driven genetic programs. (A) Length of entire hindguts in SoxC-KO E14.5 embryos and non-Cre (unrecombined SoxC fl/fl) littermates. (B) Representative immunofluorescence (IF) staining of SoxC-KO and non-Cre littermates E14.5 hindguts (blue DAPI, green Phalloidin, grey E-Cadherin). Scale bar: 50 μm. (C) Measurement of stromal widths (defined as the distance between the lower epithelial border and the beginning of the smooth muscle layer) in multiple images similar to 2B. Each dot represents one measurement, between 3 to 5 measurements were taken from n=3 SoxC-KO and n=3 WT hindguts. (D) Schematics of the multi-omics experiments performed using E14.5 wild-type (WT) and SoxC-KO hindguts. (E) UMAP plot of SoxC-KO E14.5 hindgut scRNAseq integrated with previously published E14.5 WT hindgut scRNAseq. Dots represent single cells, colored by cell type as illustrated in Figure S4E. (F) Volcano plot of gene expression fold changes in scRNAseq of E14.5 hindguts in SoxC-KO or WT mice. Magenta dots represent genes significantly downregulated in SoxC-KO (log2FC (log2-fold change) < -0.6, p_adj.glb (p value adjusted global) < 0.05), blue dots represent genes significantly upregulated in SoxC-KO (log2FC > 0.6, p_adj.glb < 0.05). Selected genes are labeled. Differential expression and p values were calculated via hierarchical permutation tests. (G) Volcano plot of ATACseq peaks in SoxC-KO and WT E14.5 hindguts as identified via the ATACseq snakePipes pipeline. Magenta dots represent peaks significantly lost or reduced in SoxC-KO compared to WT (logFC < -1, FDR < 0.05), blue dots represent peaks significantly gained or increased in SoxC-KO compared to WT (logFC > 1, FDR < 0.05). (H) ChIPseq profiles in selected genomic regions bound by Sox4 and Sox11, showing two examples of positively regulated (top) and two negatively regulated (bottom) genes, also identified in scRNAseq and ATACseq (2F&G). (I) Number of genes with significant changes detected in scRNAseq, ATACseq or ChIPseq and schematics of overlaps between the three assays. Note that the Sankey plot shows only the genes shared between the datasets, while the total number of genes in each category exceeds the number of shared genes. (J) Representative IF staining of E14.5 SoxC-KO and SoxC fl/fl hindguts (blue DAPI, grey Sox11, magenta Klf4, green Phalloidin) and quantification of nuclear Klf4 measured as averaged intensity of all nuclei in the section. (K) qPCR analysis of *Klf4* expression in WT and SoxC-KO E14.5 hindguts. (L) Representative IF staining of E14.5 SoxC-KO and SoxC fl/fl hindguts (blue DAPI, grey E-Cadherin, magenta Tead2, green Phalloidin) and quantification of nuclear Tead2 measured as averaged intensity of all nuclei in the section. (M) qPCR analysis of *Tead2* expression in WT and SoxC-KO E14.5 hindguts. (N) Representative IF staining of E14.5 SoxC-KO and SoxC fl/fl hindguts (blue DAPI, magenta Mdk, green Phalloidin) and quantification of nuclear Mdk measured as averaged intensity of all nuclei in the section. (O) qPCR analysis of *Mdk* expression in WT and SoxC-KO E14.5 hindguts. Statistical analyses for 2B and 2J-O were performed using a two-tailed unpaired t-test. Scale bars 50 μm.

To elucidate the full genetic network regulated by SoxC TFs, we conducted a multi-omics approach, integrating scRNAseq, Assay for Transposase-Accessible Chromatin sequencing (ATACseq), and Chromatin Immunoprecipitation coupled with sequencing (ChIPseq) from SoxC-KO and wild-type (WT) embryonic hindguts (Figure 2D). We identified significant gene expression changes linked to intestinal development and suggesting premature differentiation (Figures S4F-H). In SoxC-KO epithelium, we observed appearance of additional clusters and a marked upregulation of more differentiated and adult cell markers like *Lgr5, Klf4*, and *Klf5* compared to WT cells, while embryonic markers such as *Nnat, Tead2*, and *Mdk* were downregulated (Figures 2E-F, S4E-H, S5A and S5C-D). Notably, the integration of SoxC-KO scRNAseq data with previously published datasets revealed the premature appearance of adult-like cell clusters, demonstrating the role of SoxC TFs in maintaining the embryonic state (Figure S5B).

The ATACseq results highlighted substantial chromatin accessibility changes in SoxC-KO samples, reinforcing the role of SoxC TFs in modulating both gene expression and chromatin dynamics (Figure 2G). Regions with gained chromatin accessibility are associated to genes linked to differentiated intestinal cellular processes, while lost accessibilities are linked to developmental processes (Figures 2G and S6A-B). By integrating ChIPseq (Figure S6C), ATACseq, and scRNAseq data, we mapped specific genomic regions where SoxC modulates gene expression and chromatin dynamics, in direct or indirect manner. The latter is based on presence or absence of SoxC ChIPseq peaks, respectively. Noteworthy SoxC targets include *Tead2* (direct) and *Mdk* (indirect) (Figure 2H). Remarkably, the majority of differentially expressed genes were upregulated in the SoxC-KO cells. Similarly, we observed more gained chromatin accessibility than lost in the KO tissues (Figure 2I). This analysis positions SoxC as a key gatekeeper of the embryonic progenitor state, actively repressing the transition to adult stem-like states and differentiation.

Further analyses on tissues confirmed the upregulation of Klf4 and Klf5 and the downregulation of Tead2 and Mdk at both the protein and transcriptional levels (Figures 2J-O and S6D-E). Taken together, these results underscore the critical role of SoxC TFs, and their downstream components, in regulating the embryonic progenitor state during intestinal development and set the stage for investigating their role in the cancer context.

### SoxC oncoembryonic signature predicts progression of CRC

Using the intersection of omics datasets from embryos, we constructed a molecular landscape of the SoxC program, identifying genes directly or indirectly downstream of SoxC TFs, including both gene activation and repression (Figure 3A). For instance, genes such as *Smad1/5/3, Tgfbr3, Tead2, Mdk* regulate proliferation and growth; *Mif, Pten, Gldc* are involved in cell survival; *Il1r1, Il19, Rarb* play roles in immune modulation; while *Hdac9/11, Dnmt3a, Smarcd1/3* are key regulators of chromatin remodeling. Moreover, genes like *Fscn1, Cldn4, Twist1* and *Cdh1* modulate cell adhesion and migration (Figure 3A). The identified genetic profile highlights the critical role of the SoxC oncoembryonic program in orchestrating multiple fundamental cellular functions during embryonic development, and suggest that it may similarly drive CRC progression by promoting malignant behavior.

**Figure 3.**
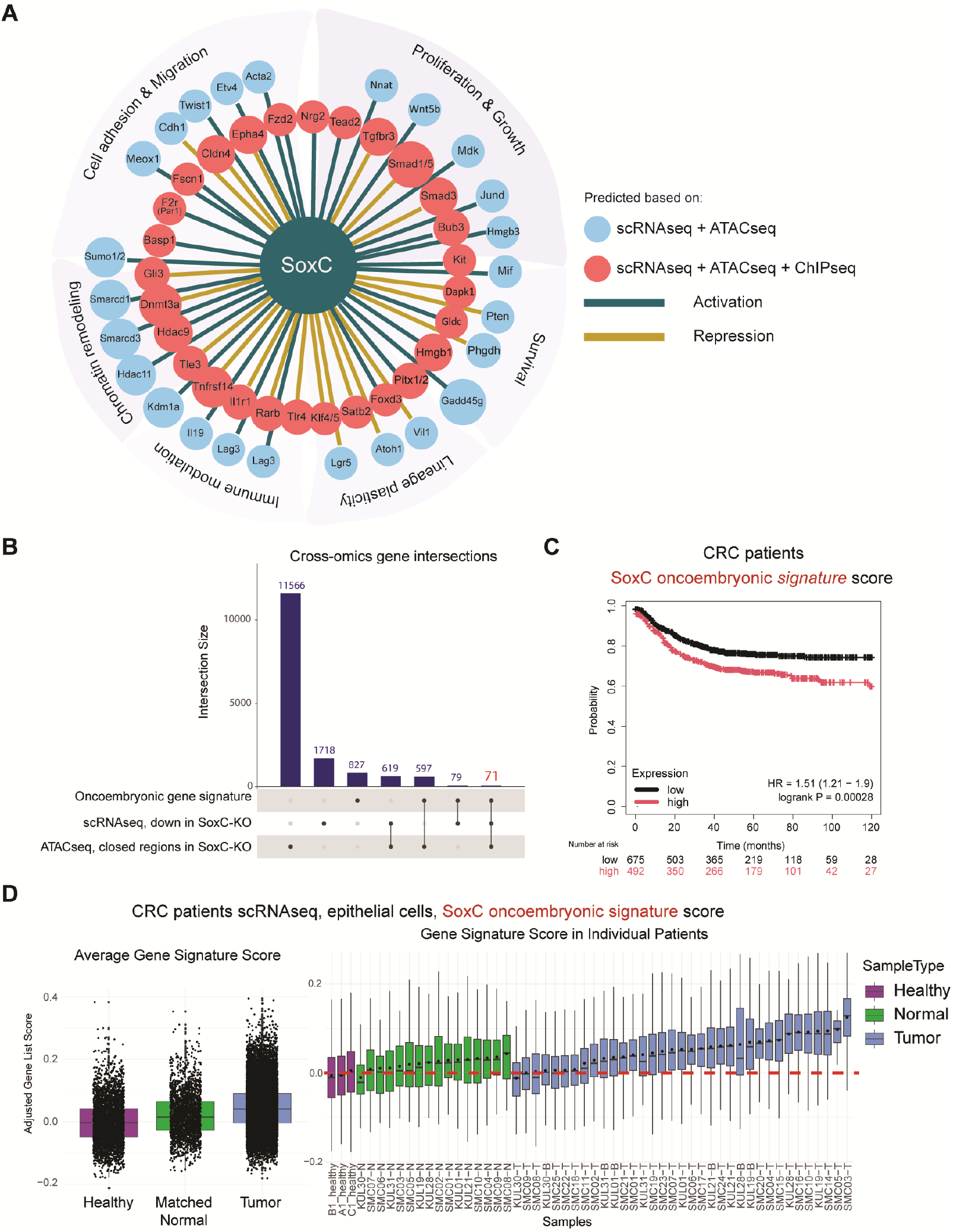
SoxC oncoembryonic signature involves several cellular functions and predicts disease outcome. (A) Predicted gene regulatory network part of SoxC genetic program, based on the integration of data from embryonic hindgut scRNAseq (Figures 2E-F), ATACseq (Figure 2G) and ChIPseq (Figure 2H) showing direct (+ ChIP) and indirect as well as activation and repression with distinct colors. Gene products are categorized based on their known functions, some with multiple functions are only shown in one category for simplicity. (B) Upset plot showing cross-omics gene intersections. (C) Kaplan Meier plots showing the inverse correlation between SoxC-oncoembryonic gene signature and CRC patients’ relapse-free survival. Hazard ratio (HR) and logarithmic ranked p Value (logrank P) were analyzed to infer the significance of the differences. Numbers below each graph represent number of patients at risk in any given time (months), black for low expression and red for high expression of each gene. The cut-off is automatically calculated based on the best performing threshold. (D) SoxC-oncoembryonic gene signature score using single-cell RNA sequencing (scRNAseq) in a cohort of 29 patients integrated with scRNAseq of three healthy individuals. Box plots highlight the average distribution of score levels across single cells in different sample groups (left), or individual samples (right). ‘Normal’ refers to phenotypically normal adjacent tissue from the same patients. The dashed red line marks the average score value of the merged three Healthy samples.

To evaluate the prognostic significance of the SoxC-regulated oncoembryonic gene expression program, we generated a SoxC-oncoembryonic gene signature including only positively regulated genes to score CRC data from patients. A 71-gene signature was derived through cross-omics gene intersection of multiple datasets, including: the oncoembryonic signature as described above (Figure 1B), genes downregulated in scRNAseq from SoxC-KO embryonic hindgut compared to WT (Figure 2F), and genes associated with chromatin regions that lost accessibility in SoxC-KO embryonic hindgut as determined by ATACseq (Figures 2G and 3B). This signature includes predicted direct targets of SoxC TFs such as *Tead2, Fscn1* (Figure 2H), *Basp1, Hdac9, Gldc*, and *F2r*. It also includes numerous indirectly regulated genes (no identified peak in ChIPseq), most notably *Mdk*, which based on predicted protein association network analysis could play a central role (Figure S7A).

We computed SoxC-oncoembryonic signature scores for patient samples and examined their correlation with disease progression. In a cohort of 1167 colon cancer patients, high average expression of the SoxC-oncoembryonic signature was significantly associated with decreased survival compared to patients with low signature expression (Figure 3C). Further analysis of scRNAseq data from 29 CRC patients (Lee et al., 2020) and three healthy colon samples (Parikh et al., 2019) revealed that the majority of tumor samples exhibited elevated expression of the SoxC-oncoembryonic signature (Figure 3D). These findings underscore the potential of the SoxC oncoembryonic signature as a prognostic biomarker and therapeutic target in CRC.

### SoxC-deficiency reduces CRC progression and prevents metastasis

To directly assess the role of SoxC TFs in tumorigenesis and metastasis, we generated SoxC-KO AKPS organoids. We simultaneously targeted the single exon of *Sox4, Sox11*, and *Sox12* at two distinct sites each within their coding regions with enhanced Cas12a mediated perturbations (Figure 4A)(Campa et al., 2019; DeWeirdt et al., 2021). Control AKPS organoids, in which intergenic regions were targeted using the same method with the same number of perturbations, displayed behavior akin to parental organoids: all injections led to primary tumor engraftment, with some animals developing macroscopic liver metastases. In contrast, SoxC-KO tumors displayed significantly reduced tumor size, and none of the animals exhibited detectable liver metastases, highlighting the critical role of SoxC in both tumor growth and metastatic spread (Figures 4B-D).

**Figure 4.**
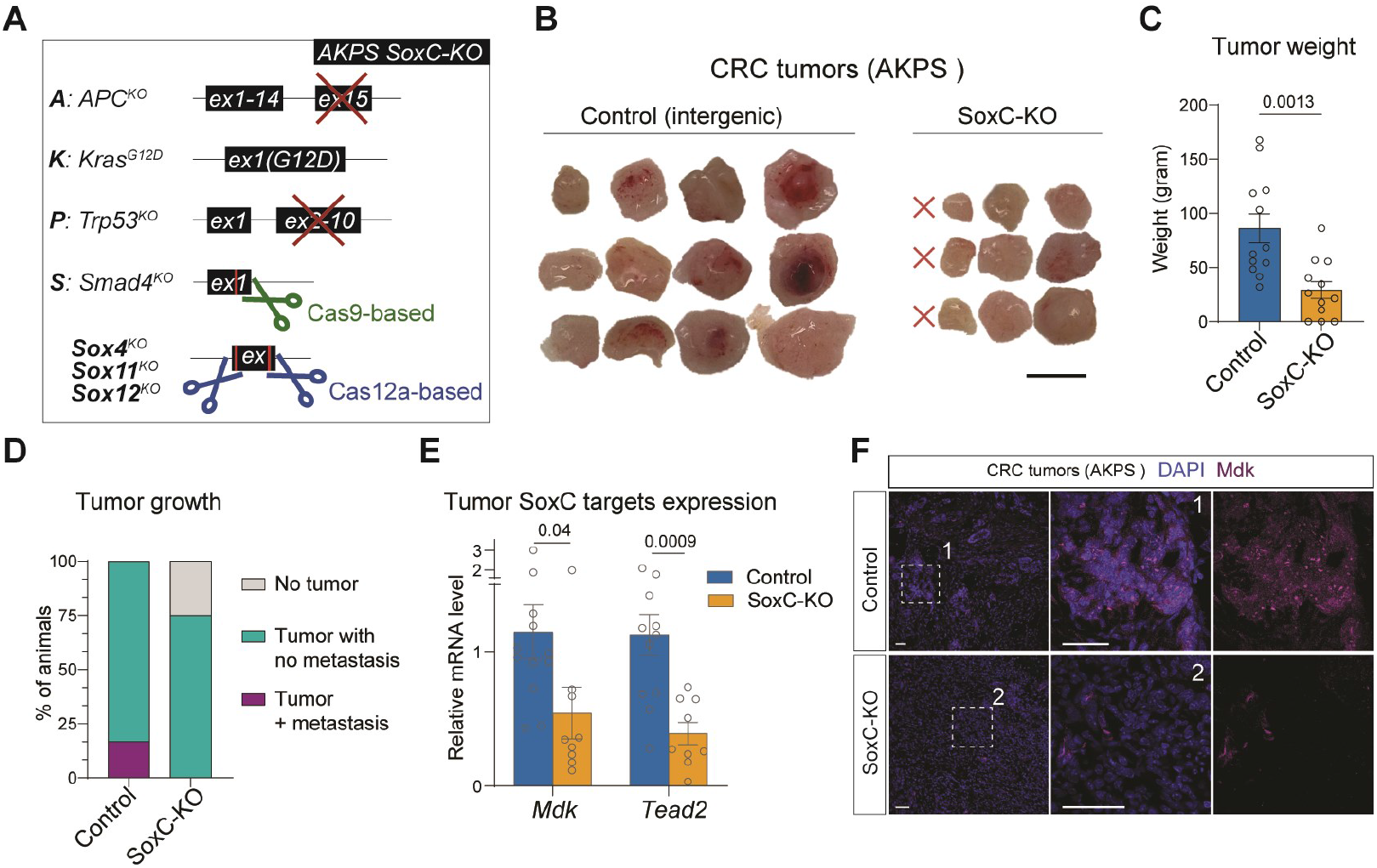
SoxC KO reduces tumor growth and blocks metastasis. (A) Schematics of the generation of AKPS SoxC-KO organoids. (B) Primary tumor morphology of control (Cas12a intergenic targeting) and SoxC-KO (Cas12a targeting *Sox4, Sox11* and *Sox12* each with two specific gRNAs). Scalebar: 5 mm. (C) Bar plot of the tumor weights in the two groups. (D) Percentage of primary tumor and metastatic growth. (E) Quantitative PCR showing the transcript levels of SoxC TFs targets, *Mdk* and *Tead2*. (F) Immunofluorescent staining of Mdk in Control and SoxC-KO colorectal cancer (CRC) tumors. Scalebars: 50 μm. Statistical analyses were performed using an unpaired t-test.

Bulk transcript analysis of SoxC-KO tumors confirmed reduced expression of SoxC TFs and downregulation of key targets such as *Tead2* and *Mdk* (Figures 4E-F and S7B). These findings suggest that SoxC governs a genetic program critical for maintaining tumor growth and facilitating metastasis, further solidifying SoxC as a promising therapeutic target.

### Tead2 and Mdk are critical SoxC downstream factors

The YAP-TAZ cofactor Tead2 and the embryonic growth factor Mdk emerged as key SoxC-regulated oncoembryonic genes in multiple analyses (Figures 1B, 2F-H, 2J-O and 4E-F). We therefore wanted to further explore their role in SoxC program. *In situ* protein localization analysis showed that both Mdk and Tead2 share the same expression pattern as SoxC TFs: high expression in the embryonic hindgut, downregulated in healthy adult tissue, and reactivated in CRC (Figures 5A-B).

**Figure 5.**
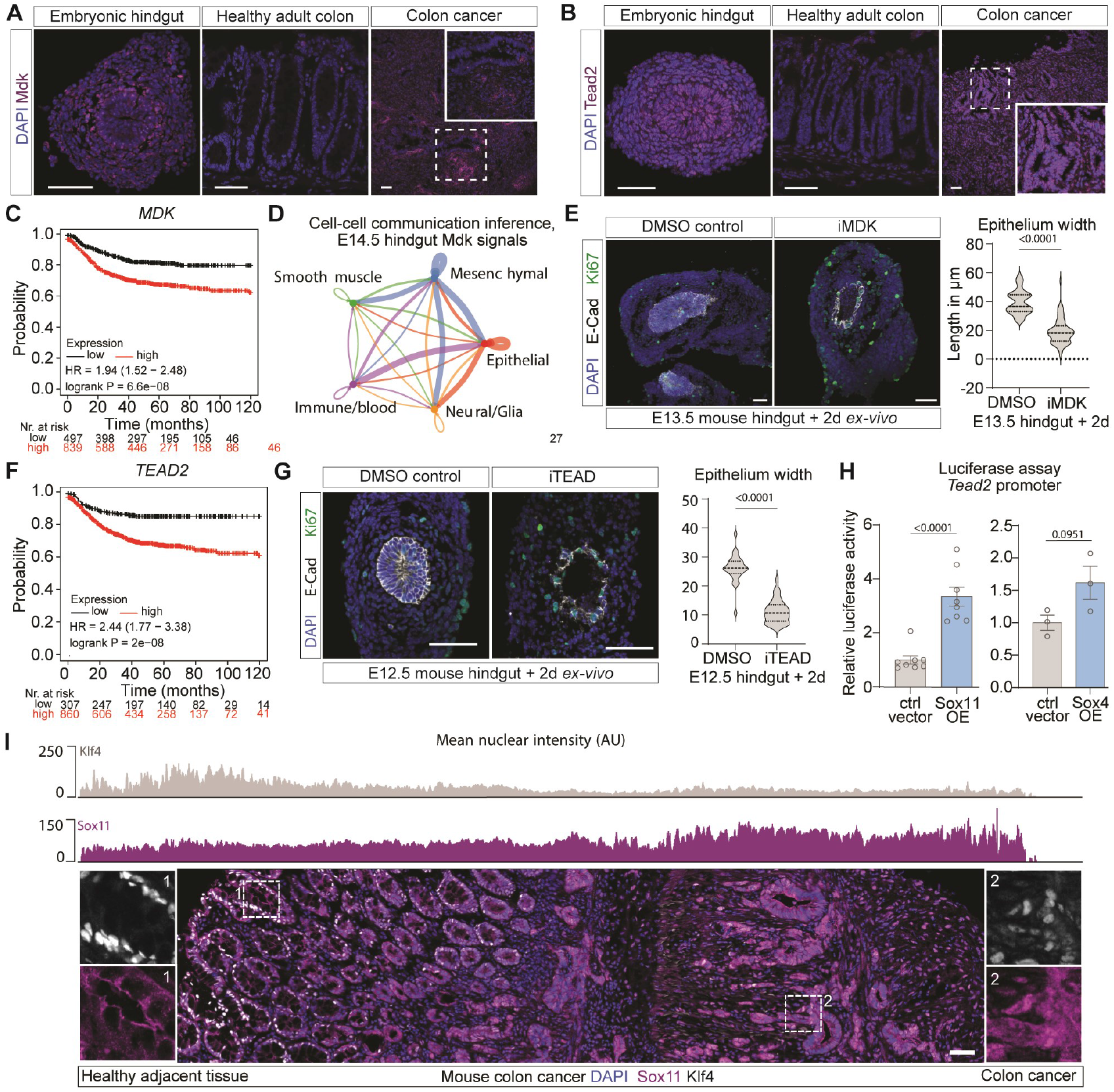
Tead2 and Mdk are key downstream targets of SoxC TFs. (A) Immunofluorescent (IF) staining of Mdk (magenta) in E14.5 hindgut, healthy adult colon and colon cancer. Blue DAPI. (B) IF staining of Tead2 (magenta) in murine E14.5 hindgut, healthy adult colon and colon cancer. Blue DAPI. (C) Survival analysis of colorectal patients divided into high or low *MDK* expression. (D) Inferred interactions through Mdk signaling in E14.5 WT hindgut. Edge weights are proportional to the interaction strength. (E) IF staining and quantification of epithelial thickness of E13.5 hindgut explants cultured *ex-vivo* for 2 days in the presence of MDK inhibitor (iMDK, n=6) or control (DMSO, n=6). For each explant, 5-7 measurements were taken. Blue DAPI, grey E-cadherin, green Ki67. (F) Survival analysis of colorectal patients divided into high or low *TEAD2* expression. (G) IF staining and quantification of epithelial thickness of E12.5 hindgut explants cultured *ex-vivo* for 2 days in the presence of TEAD inhibitor (iTEAD, n=4) or control (DMSO, n=3). For each explant, 5-7 measurements were taken. Blue DAPI, grey E-cadherin, green Ki67. (H) Activity of *Tead2* promoter after Sox11 or Sox4 transfection assessed by luciferase assay in HEK293 cells. N=8 for Sox11, n=3 for Sox4. (I) IF staining of Klf4 and Sox11 in mouse colon cancer and adjacent tissue. Insets of representative regions with high nuclear Klf4 or Sox11 expression. On top: quantification of mean nuclear intensity along a left-to-right axis. Scale bars: 50 μm. Statistical analyses for (D), (G), (H) using two-tailed unpaired t-test.

Consistent with the worse survival associated with high SoxC signature expression, elevated *MDK* levels were also correlated with poor survival in CRC patients (Figure 5C). Cell-cell communication analysis suggests Mdk functions as a secreted ligand during hindgut development (Figures 5D and S8A-B). In line with this, inhibition of Mdk in hindgut explants resulted in epithelial flattening and impaired development (Figure 5E). As a secreted ligand, Mdk likely influences various cell populations in the tumor microenvironment, particularly immune cells as evidenced by our functional genetic network and cell-cell communication analysis (Figure S2D and S8C-D).

Similarly, high *TEAD2* expression correlated with poor survival in CRC patients (Figure 5F). Inhibition of Tead activity in hindgut explants led to impaired epithelial development, producing a flattened epithelium, like the effects observed with Mdk inhibition (Figure 5G). Additionally, Tead inhibition reduced the number of proliferating cells (Figures S8E-F), further linking SoxC regulation of *Tead2* to cellular proliferation.

To further explore the regulation of *Mdk* and *Tead2* by SoxC TFs, we analyzed their modes of regulation. Our ChIPseq data (Figure 2H) predicted direct regulation of *Tead2* by both Sox11 and Sox4, which we confirmed through reporter assays (Figures 5H and S8G). To identify potential intermediate regulators of indirect targets, e.g. *Mdk*, we analyzed chromatin regions that were differentially accessible in SoxC-KO versus WT samples, and the promoter of genes with differentially accessible regions. By filtering based on differential expression in SoxC-KO versus WT, as well as embryo versus adult tissues scRNAseq, we identified Hoxa13, Elk4, Hnf4a, Tcf3, Esrra, and Klf3/4/5/6 as potential intermediaries (Figure S8H). Except for Tcf3, all these transcription factors were upregulated in SoxC-KO compared to WT hindgut and adult cells compared to the embryo (Figures S8I-J). This suggests a mechanism where these TFs, repressed by SoxC in the hindgut, act to suppress embryonic programs in adult tissues. This hypothesis was supported by the inverse correlation of protein localization between Klf4 and Sox11 in mouse CRC samples: healthy adjacent colon regions showed high Klf4 but low Sox11, while cancerous lesions exhibited high Sox11 and low Klf4 (Figures 5I and S8K).

Overall, these results confirm the importance of SoxC-regulated Tead2 and Mdk in hindgut development and suggest a mechanism of reactivation in CRC, highlighting their potential roles in tumor progression.

## Discussion

Reawakened embryonic programs have long been assumed to contribute to cancer phenotypic plasticity (Virchow, 1859; Krebs, 1947; Gold and Freedman, 1965; Weinberg, 1996; Nieto et al., 1994; Hanahan and Weinberg, 2011; Nieto, 2013; Aiello and Stanger, 2016; Hanahan, 2022; Fazilaty and Basler, 2023; Fazilaty, 2023), yet how these programs would promote tumor progression has remained elusive. In this study, we applied a new approach— *oncoembryology*—to identify and dissect the reactivated embryonic programs in CRC. To fully harness the potential of oncoembryology, we compared single-cell transcriptomics from hindgut development, CRC tumor progression, and healthy adult colon tissues. SoxC TFs emerged as regulators of both embryonic development and CRC. Returning to the embryonic context allowed us to define the core components of the SoxC program using conditional loss-of-function models and a multi-omics approach. Cross-system comparisons then enabled us to pinpoint *bona fide* downstream targets of SoxC, which were then validated through functional assays. While the Sox family has been implicated in intestinal development, the specific role of the SoxC subfamily has not been previously established (Fu and Shi, 2017). SoxC TFs, particularly Sox4 and Sox11 are known regulators of neural development and early organogenesis (Bergsland et al., 2006, p. 11; Bhattaram et al., 2010). Our findings demonstrate that SoxC TFs are master regulators in maintaining the embryonic progenitor state during intestinal development as well as driving cancer progression. Although SoxC factors were previously linked to wound healing and other cancer types including breast and prostate (Miao et al., 2019; Oliemuller et al., 2020), their critical role in colorectal cancer has remained underexplored.

We found that SoxC factors potentially regulate a range of essential cellular processes, including proliferation and growth, cell survival, adhesion and migration, lineage plasticity and immune modulation. Several predicted direct targets of SoxC in the embryo are well-established regulators of these key functions: Basp1 (involved in neuronal development and plasticity), Hdac9 (a histone deacetylase linked to chromatin remodeling and immune regulation in cancer)(Yang et al., 2021), Gldc (critical for cancer cell survival and stemness)(Zhang et al., 2012), and F2r (thrombin receptor PAR1, implicated in tumor growth and metastasis)(Hua et al., 2021). A key finding of our study is the observation that SoxC TFs negatively regulate many gene expression programs. For example, we found that Klf4, a transcription factor crucial for maintaining epithelial homeostasis in the intestine (Hickey et al., 2023; Katz et al., 2002), was upregulated in SoxC-KO.

Loss of function of the SoxC TFs in the embryo, therefore, disrupted the balance between progenitor maintenance and differentiation, leading to premature differentiation of epithelial cells. This regulatory mechanism likely contributes to tumor progression in CRC. Given the established role of Klf4 in repressing epithelial-to-mesenchymal transition and maintaining epithelial integrity (Agbo et al., 2019; Borrelli et al., 2024; Liu et al., 2012), its inhibition by SoxC may further drive tumor aggressiveness by promoting a loss of epithelial cohesion.

We also looked at the loss of function of the SoxC TFs in tumors: the tumors were smaller and had reduced metastatic potential. Indeed, we failed to find any liver metastases in SoxC-KO models. Supporting analyses demonstrate that the SoxC oncoembryonic gene signature, derived from our integrative analyses is a robust predictor of poor survival in CRC patients. These results highlight the therapeutic potential of targeting SoxC programs. Two downstream targets Tead2 and Mdk are of particular interest.

Targeting Mdk and Tead factors has been explored as a therapeutic strategy in other tumor systems. Mdk is known to promote an immune-evasive microenvironment in melanoma, leading to resistance to immune checkpoint blockade and poor makes tumors more sensitive to immunotherapy. (Cerezo-Wallis et al., 2020). MDK has also been implicated in immune tolerance in CRC, correlating prognosis. However, genetically targeting Mdk makes tumors more sensitive to immunotherapy. (Cerezo-Wallis et al., 2020). MDK has also been implicated in immune tolerance in CRC, correlating with poor patient survival (Hashimoto et al., 2024). Our study shows that Mdk is a key signaling component in CRC that cancer cells could use to influence different cell types in the tumor microenvironment, highlighting the potential benefits of blocking it.

Inhibition of TEAD factors has been explored in several cancer contexts. Combining TEAD inhibitors with targeted therapies, such as EGFR or MEK inhibitors, has shown synergistic effects in preclinical models. Multi-component inhibition may offer more effective therapeutic benefits by tackling both drug resistance mechanisms and multiple cancer-causing factors at the same time (Pobbati et al., 2023). A more comprehensive understanding of the molecular program, particularly the oncoembryonic factors and their interactions, is essential for identifying vulnerabilities and enabling the development of more effective combinatorial therapies. Targeting oncoembryonic factors, in particular, offers the potential for disease-specific treatments, sparing healthy cells that do not express these factors.

Taken together, our results highlight the critical role of the SoxC oncoembryonic program in regulating multiple cellular functions during embryonic development and suggest that this program similarly drives malignant behavior in colorectal cancer. Oncoembryology not only bridges the gap between developmental biology and cancer research but also lays the groundwork for future studies aimed at exploiting these shared pathways for clinical benefit.

## Materials and Methods

### Patient samples

Written informed consent was obtained before specimen collection, and the study was approved by the local ethics committees (Cantonal Ethics Committee of the Canton Zurich, approval no. EK-1755). Three tumor samples were resected from colorectal cancer patients. Two healthy colon specimens were resected from heathy colon regions. After surgical resection, tissue samples were fixed in 4% formaldehyde for 2 hours and incubated in 30% sucrose overnight at 4°C before cryosectioning (see below).

### Mice

C57BL/6J wild-type mice were purchased from Charles River (Strain code: 027), C57BL/6-NRj were purchased from Janvier, Rosa26-CreERT2 (R26-CreERT2) mice were purchased from The Jackson Laboratory (Strain code: 008463), Sox4 fl/fl; Sox11 fl/fl; Sox12 fl/fl (SoxC fl/fl, (Bhattaram et al., 2010; Hoser et al., 2008; Miao et al., 2019)) mice were a gift from Véronique Lefevbre (Children’s Hospital of Philadelphia, USA) and Elizabeth Sock (Institute of Biochemistry, FAU Erlangen-Nuernberg, Germany). SoxC fl/fl mice were bred to R26-CreERT2 and genotyped using the primers as in Table S1. Standard PCR reactions were mixed. Sox12 PCR required the addition of 5% DMSO. For Sox11 and Sox4 a standard three-step PCR was performed as follows: 2 min denaturation at 94°C, 35 cycles of 15 sec at 94°C, 75 sec at 65°C, 90 sec at 72°C, 2 min final elongation at 72°C. Sox12 and R26-CreERT2 were performed as follows: 2 min denaturation at 94°C, 35 cycles of 15 sec at 94°C, 30 sec at 57°C, 45 sec at 72°C, 2 min final elongation at 72°C. We affirm compliance with all relevant ethical regulations for animal testing and research. All mouse experiments were performed in accordance with Swiss Guidelines and approved by the Cantonal Veterinary Office Zurich, Switzerland. The mice were all housed in the same accredited facility under specific-pathogen free conditions in a 12 h-12 h light-dark schedule, and chow and water were provided ad libitum. At the experimental end-point, mice were euthanized by CO2.

### Timed breeding

A C57BL/6J male and a C57BL/6J female or a R26-CreERT2; SoxC fl/fl male (for RNA extraction) or SoxC fl/wt male (for immunofluorescent stainings) and a SoxC fl/fl female at age between 7-24 weeks were transferred to the same cage in the evening. In the morning the female was checked for the presence of a vaginal plug. If present, the day was counted as embryonic day E0.5. To induce SoxC knock-out, gavage was used to administer tamoxifen dissolved in corn oil to a final dose of 200 mg per kg of body weight.

### Colonoscopy-guided submucosal injection of CRC organoids

This procedure was adapted from a previously published protocol (Roper et al., 2018). 36 hours after passaging, AKPS organoids were mechanically dissociated and resuspended in OptiMEM (Gibco). Each 40uL Matrigel® (Corning) dome was resuspended in 50 μL OptiMEM for one injection in one mouse. Male C57BL/6NRj mice between the ages of 7 and 16 weeks were anesthetized using isoflurane inhalation and placed on a heating pad at 37°C facing upwards. The colon was rinsed with pre-warmed PBS to remove feces using the plastic tubing from an intravascular catheter (BD) mounted onto a 50 ml syringe (B. Braun). The organoid solution was injected with custom injection needles (33 gauge, 400 mm length, point style 4, 45° angle, Hamilton), a syringe (Hamilton) and a colonoscope with an integrated working channel (Storz). Pressure was applied on the needle just enough to allow the delivery of 50 μL of organoid suspension below the mucosa and the formation of a visible injection bubble. Mice were monitored until an experimental or humane end point was reached.

### *Ex-vivo* culture of hindgut

Embryonic hindguts from C57BL/6J wild-type were isolated at E12.5 or E13.5 stages. Samples were divided into groups of cultured control which were treated with DMSO or treated with iMDK (Sigma, 5mg/ml) or iTEAD (TM2 TEAD inhibitor, 10μM biotechne) inhibitors. The *ex-vivo* cultures were carried out as previously described (Fazilaty et al., 2021). Briefly, the hindguts were isolated, placed on transwell plates (24 mm inserts, 8.0 mm polycarbonate membranes, Corning Coaster), and incubated for 48 hours at 37°C in BGJb medium supplemented with Ascorbic acid (0.1 μg/ml, Sigma) and antibiotics (PenStrep (10,000U/ml Penicillin; 10,000 mg/ml Streptomycin), Gibco). Samples were then fixed for 1 hour and processed as mentioned below before sectioning and immunofluorescence analyses.

### AKPS organoids

#### Culturing

*Vil-creER*^*T2*^;*APC*^*KO*^;*Trp53*^*KO*^;*Kras*^*G12D/WT*^*Smad4*^*KO*^ (AKPS) organoids were cultured in 50 μl Matrigel domes (Corning) as described previously (Sato and Clevers, 2013). To make complete medium, Advanced DMEM/F12 (Life Technologies) was supplemented with 10 mM HEPES (Gibco), 1X GlutaMax (Gibco), 1% penicillin–streptomycin (Gibco), 1X B27 supplement (Gibco), 1X N2 supplement (Gibco) and 1 mM N-acetylcysteine (Sigma-Aldrich). Organoids were split every 2-3 days by mechanical dissociation.

#### RNP-mediated Smad4 KO

*Vil-creER*^*T2*^;*APC*^*KO*^;*Trp53*^*KO*^;*Kras*^*G12D/WT*^ (AKP, Figure S1A) organoids, a gift from Owen Sansom (Beatson Institute for Cancer Research), were cultured under the above-described conditions with supplementation of 100 ng/mL mouse recombinant noggin (Sigma). Two days before transfection, 5 nM nicotinamide (Sigma) and 10 μM ROCK inhibitor (Y-27632, Stem Cell Technologies) were added to the medium. sgRNA with previously published sequence (de Sousa e Melo et al., 2017)(Figure S1A) were obtained from IDT as a combination of both crRNA and tracrRNA (Alt-R system). Organoids were collected, washed, resuspended in TrypLE (Gibco) and dissociated into single cells by incubating at 37 °C for 10 min. After neutralization with Advanced DMEM/F12, cells were centrifuged at 290 x g for 3 min, resuspended in 1 mL complete medium with 5 mM nicotinamide and 10 µM ROCK inhibitor and transferred to two wells of a 48-well plate containing 50 µL of transfection mix. The transfection mix was prepared by combining 25 μL OptiMEM containing 1,250 ng Alt-R S.p. Cas9 Nuclease V3 (Integrated DNA Technologies), 240 ng Mm Cas9 SMAD 4.1.AR Alt-R CRISPR-Cas9 sgRNA (Integrated DNA Technologies) and 2.5 μL Cas9 Reagent Plus (Invitrogen) with 25 µL OptiMEM containing 1.5 μL CRISPRmax reagent (Invitrogen) according to manufacturer’s instruction (CRISPRmax Cas9 Transfection Reagent, Invitrogen). sgRNA was omitted in the control condition. The cells were spinoculated for 1 h by centrifuging at 600 x g and 32 °C, then incubated for 2 h at 37 °C. Cells were then collected, washed once with Advanced DMEM/F12, resuspended in Matrigel and cultured as described above (with supplementation of 10 µM ROCK inhibitor in the first passages). After three days, 50 ng/mL TGFβ (Peprotech) was added to the medium of both CRISPR-edited and control organoids for metabolic selection and kept for two further passages (Figure S1C). Smad4 genomic locus was amplified by PCR (forward primer 5’-ATCTACCTTGTGAAATGTGTTCTC-3’, reverse primer 5-TACCAAACTCTCAATTGCTC-3’) and the presence of small insertions and deletions assessed with the Guide-it Mutation Detection Kit (Takara Bio). The PCR amplicons were then inserted into pGEM vectors using the pGEM-T Easy Vector system (Promega) and 40 (CRISPR-edited) and 16 (control) bacterial colonies were sequenced by Ecoli NightSeq (Microsynth). Sequencing results were then aligned to a reference sequence in CLC Main Workbench 7 (Qiagen) to assess their mutational status (Figure aS1D).

#### enAsCas12a-based knock-out of SoxC

AKPS SoxC knockout organoids were generated using enAsCas12a (DeWeirdt et al., 2021; Kleinstiver et al., 2019) in combination with Pol II delivered crRNA-arrays (Campa et al., 2019). The necessary targeting sequences for *Sox4, Sox11, Sox12* and intergenic regions, were obtained using the guide design tool CRISPick (DeWeirdt et al., 2021; Kim et al., 2018) with the mouse GRCm38 reference genome and enAsCas12a CRISPRko mode. The *Sox4, Sox11*, and *Sox12* crRNAs (spacer – direct repeat) were combined in sets of two to form a 6-mer pre-crRNA array (AKPS-SoxCKO) with efficacy scores > 0.8 for each guide. Similarly, 6-mer pre-crRNA arrays were designed for intergenic regions perturbing the SoxC chromosomes with two crRNAs per chromosome. The arrays were cloned into a 3^rd^ generation lentiviral vector which expressed the array under the EF1a promoter together with GFP and Puromycin (EF1a-EGFP-2A-Puro-Triplex-pre-crRNA array-WPRE)(Campa et al., 2019) to be compatible with enAsCas12a-2A-Blas (pRDA_174, Addgene, 1346476) (DeWeirdt et al., 2021) in AKPS organoids. For lentiviral production, 3 million HEK cells were seeded in a 75 cm2 flask in DMEM (gibco) + 10% heat-inactivated FBS (gibco) + 1% penicillin– streptomycin (gibco). After 24 h cells were transfected using Lipofectamine™ 3000 reagent (ThermoFisher) according to manufacturer’s instruction. 2.25 μg pRev (Addgene 12253), 2.25 μg pVSV (Addgene 12259) and 6.75 μg pMDL (Addgene 12251) plasmids were mixed with 9 μg of lentiviral expression vector, 500 μL OptiMEM and 40.5 μL P3000 reagent. 28 μL Lipo3000 were added to 500 μL OptiMEM, mixed to the DNA mixture and added to the cells. After 8 h, the medium was collected and stored at 4°C and replaced by lentivirus packaging medium (5% FBS + 0.2 mM Sodium Pyruvate in OptiMEM). After 48 h, the medium was collected, pooled to the previously collected medium, filtered through a 0.45 μm filter and concentrated using an Amicon® Ultra-15 Centrifugal Filter Unit (Merck Millipore) to 1 mL and subsequently stored at -70°C. For lentiviral transduction, organoids were dissociated mechanically, incubated in TrypLE for 5 min at 37°C and washed with Advanced DMEM/F12 + 10% heat inactivated FBS. 200 μL of concentrated virus were used to transduce organoids from 4-6 40 μL Matrigel domes. Complete medium (as described above) + 4 ng/μL Polybrene + 10 μM ROCK inhibitor + 5 mM nicotinamide (Sigma) was added up to 500 μL and cells were spinoculated for 1 hour 600 x g at 32°C. Cells were then let to recover at 37°C for 2-4 hours before being plated in Matrigel and cultured in complete medium + Y-27632 + nicotinamide. After outgrowth, organoids were selected for 14 days using 2 μg/mL of Puromycin (Sigma) and 8 μg/mL of Blasticidin (ThermoFisher).

### Single-cell RNA sequencing

#### Single-cell isolation and preparation for scRNAseq of tumors

Mice were euthanized by CO_2_ and the abdomen opened. The colon was extracted and examined for the presence of tumors. The intestinal tube was cut open and rinsed with PBS (gibco). Healthy-looking tissue was removed from around the tumor with a blade. Tumors from three mice (replicate 1) and three mice (replicate 2) were used for subsequent steps. Tumors were minced in small pieces and subsequently dissociated into single cells using the Tumor Dissociation Kit (Milteny Biotec) as per manufacturer’s instructions using the gentleMACS™ Octo Dissociator with Heaters. DNase I (Roche) at a final concentration of 25 μg/mL was added to all washing steps to remove DNA and reduce viscosity. Single cells from each tumor were stained for antibodies against the epithelial marker Epcam (Invitrogen) and the immune marker CD45 in 1mL of staining solution (PBS (gibco) + 0.2% BSA (Roche)

+ 2 μL Epcam-PE-Cy5 (eBioscience) + 2 μL CD45-APC-eFluor780 (eBioscience). Cells were then centrifuged 7 min at 300 x g and resuspended in 200 μL ADMEM/F12 (Gibco) + Hepes (Gibco, 100mM) + Glutamax (Gibco, 1X) + 0.2 μL DAPI (ThermoFisher). Epithelial, immune and double negative cells were sorted from each sample using a BD FACS Aria III 4L operated by the Cytometry Facility at UZH. Single stained samples were used for compensation. For each mouse cells were mixed in the following ratios: 35% epithelial cells, 5% immune cells, and 60% double-negative cells. Approximately 60K cells for each replicate with a similar proportion of cells from each mouse were used for library generation.

#### Single-cell isolation and preparation for scRNAseq of E14.5 SoxC-KO hindguts

Two pregnant females were euthanized by CO_2_ and the abdomen opened and the uteri extracted. Amniotic sacs were opened to retrieve the embryos. Hindguts were dissected and collected in BGJb medium (gibco) + ascorbic acid (0.1 μg/mL, Sigma). A total of 19 hindguts were minced with a blade and incubated for 30 min at 37°C in a solution of Collagenase D (Roche, 1 mg/mL). After filtering in a 40 μM strainer and washing with BGJb, the single cells of each litter were stained for the epithelial marker Epcam in 500 μL staining solution (PBS (gibco) + 0.2% BSA (Roche) + 1 μL Epcam-PE-Cy5 (eBioscience)). For each litter, cells were mixed at a ratio of 2:1 negative to epithelial cells. A total of 50K cells were used for library generation.

#### Generation of scRNAseq libraries

Whole transcriptome analysis (WTA) libraries were generated following BD Rhapsody standard protocols (Doc ID: 210966, Doc ID: 23-21711-00) using the following kits: BD Rhapsody™ Enhanced Cartridge Reagent Kit (BD 664887); BD Rhapsody™ Cartridge Kit (BD 633733); BD RhapsodyT™ cDNA Kit (BD 633773); BD Rhapsody™ WTA Amplification Kit (BD 633801). Libraries were indexed using different Library Reverse Primers (650000080, 650000091-93) for each sample.

#### Sequencing of libraries

Libraries were sequenced at the Functional Genomics Center Zurich (FGCZ) using a single lane of a 10B flowcell on the Illumina Novaseq X Plus and 150 bp for R1/R2. A 3% PhiX spike-in was used for all libraries.

#### Mapping of data

An indexed genome was generated with STAR (Dobin et al., 2013) (version 2.7.10b) using the following reference files: GRCm38.p6 and GENCODE’s M25 annotation. Raw .fastq were aligned and unique alignments were counted using STARsolo (Kaminow et al., 2021) (version 2.7.10b), using similar parameters as previously described (Mallona and Robinson, 2024).

### Computational analysis for scRNAseq

#### Pre-processing

For the healthy adult colon, the following published datasets were used: GSE151257 (Brügger et al., 2020) and GSE266161 (Moro et al., 2024). For embryonic E14.5, E15.5 and E18.5 WT hindgut, the following published dataset was used: GSE15400 (Fazilaty et al., 2021). For earlier embryonic hindgut, the following published dataset was used: GSE186525 (Zhao et al., 2022). Data analysis of scRNAseq was performed using the Seurat package version 5.1 (Butler et al., 2018) in R version 4.4.1. Cells with counts for less than 200 genes and genes that were detected in less than 3 cells were removed. Quality control metrics (mitochondrial transcripts, total number of genes and total number of transcripts) were calculated using addPerCellQCMetric() from the *scuttle* package and filtered using isOutlier() from the *scater* package for the number of genes and transcripts. Cells with high mitochondrial gene content were determined visually from the distribution of mitochondrial transcripts content and removed (GSE266161 healthy adult 60%, GSE151257 healthy adult 25%, AKPS-1 55%, AKPS-2 25%, GSE15400 WT embryo 25%, SoxC-KO embryo 30%, GSE186525 embryonic intestine 30%). Doublets were identified and removed using scDblFinder(). After pre-processing, the following cell numbers were retained: 4,024 (GSE266161 healthy adult), 6,120 (GSE151257 healthy adult), 5,008 AKPS-1, 1,199 AKPS-2, 5,124 (GSE15400 WT embryo), 3,738 SoxC-KO embryo, 65,460 GSE186525 embryonic intestine. For all datasets, normalization, scaling, variable gene selection and principal component analysis was performed using the standard workflow in the Seurat package (Butler et al., 2018; Stuart et al., 2019) (https://satijalab.org/seurat/).

*Dimensionality reduction, clustering and integration* Dimensionality reduction was performed using UMAP (McInnes et al., 2018) with 25 principal components (30 for oncoembryonic integration) and a resolution of 0.5 (0.8 in healthy adults integrated with tumors). Clusters were then identified using the FindNeighbors() and FindClusters() functions in Seurat, and annotated based on known epithelial and stromal marker genes (Figure S2A, S4E). For the integration of embryonic, healthy adult and tumor epithelium, clusters positive for Epcam were subset. For the integration of multiple developmental stages, (GSE186525) only cells were subset that had one of the following meta data “sample”: e95, e105_i, e115_li, e135_li, e155_l. Data integration was done using FindIntegrationAnchors() (using 50 principal components for oncoembryonic integration, 30 for all other) and IntegrateData().

#### Pseudotime, regulatory network, gene ontology, differential expression, cell-cell communication analyses

Pseudotime models for time course experiments were built using the R package *psupertime* v0.2.6 (Macnair et al., 2022). We used the package following the standard workflow using the penalization ‘best’. Regulatory network inference was performed using the R package SCENIC v1.3.1 following the standard workflow (Aibar et al., 2017). GENIE3 (Huynh-Thu et al., 2010) was used to infer the co-expression network. Transcription factor motif analysis was performed using *Rcistarget* v1.24 and the following databases: mm9-500bp-upstream-7species.mc9nr.genes_vs_motifs.rankings.feather, mm9-tss-centered-10kb-7species.mc9nr.genes_vs_motifs.rankings.feather. For cell state identification, *AUCell* v1.26.0 was used. Gene ontology and predicted genetic networks were generated using ClueGO (Bindea et al., 2009) v2.5.10 via Cytoscape v3.10.1 (Shannon et al., 2003). The highly expressed genes in cell clusters were used for the analyses. The network clustering contained data from GO_BiologicalProcess-EBI-UniProt-GOA-ACAP-ARAP-25.05.2022 and

GO_MolecularFunctions-EBI-UniProt-GOA-ACAP-ARAP-25.05.2022. Clusters (containing at least three nodes) were identified and network specificity was adjusted based on the number of genes, which originated from highly expressed genes in each cluster analyzed via the FindMarkers() function in *Seurat*. Differential expression analysis was performed using the R package *distinct* v1.16 (Tiberi et al., 2023) with default permutation numbers using log counts as input and generating log2 fold-changes using the cpm() function from *edgeR* v4.2.1 (Robinson et al., 2010). Before differential analysis, all epithelial annotated clusters were merged into one cluster, as well as all mesenchymal clusters and immune clusters. Genes were considered differentially expressed if p_adj.glb < 0.05 and log2-fold change > 0.6. Plots were generated using *ggplot2* v3.5.1 (Wickham, 2016). For cell-cell communication analysis, *CellChat* v2.1.2 (Jin et al., 2023, 2021) was used following the standard workflow. For comparison of cell-cell communication between embryonic hindgut and adult colon, healthy adult from GSE266161 was used. For cell-cell communication analysis in cancer, AKPS-1 and AKPS-2 integrated were used.

### ATACseq

17 wildtype and 14 SoxC-KO E14.5 hindguts were dissected, divided into 3 replicates (WT: 6+5+6, KO: 5+5+4) and dissociated into single cells as described above. Cells were resuspended in 100 μL PBS and counted. 50K cells of each sample were processed independently for ATACseq library preparation (Buenrostro et al., 2015). Briefly, cells were centrifuged for 5 min at 500 x g and resuspended in cold lysis buffer (10 mM Tris-HCl, pH 7.4, 10 mM NaCl, 3 mM MgCl2, 0.1% IGEPAL CA-630). After lysis, the nuclei were then centrifuged for 10 min at 500 x g, resuspended in 50 μL 1X transposition mix (25 μL Tagment DNA Buffer, 2.5 μL Tagment DNA Enzyme, 22.5 μL nuclease-free H2O) (Illumina) and incubated for tagmentation at 37°C shaking at 400 rpm for 30 min. Tagmented DNA was purified using MinElute PCR Purification Kit (Qiagen) and eluted in 10 μL elution buffer. The library was then amplified for 5 cycles using the NEBnext high fidelity 2X PCR Master Mix (NEB) and Nextera compatible Multiplex primers (ActiveMotif) in the following reaction: 10 μL transposed DNA, 10 μL water, 2.5 μL 25 μM Nextera primer 1, 2.5 μL 25 μM Nextera primer 2, 25 μL L 2X PCR Master Mix. 5 μL of the PCR reaction were used for qPCR to determine the additional number of PCR cycles to avoid saturation o the amplification. Between 5 and 8 additional PCR cycles were performed. Amplified libraries were then purified using MinElute PCR Purification Kit (Qiagen) and eluted in 20 μL elution buffer. DNA concentration was quantified at a Qubit (Thermofisher). Libraries were sequenced by the FGCZ using a single lane of a 10B flowcell on the Illumina Novaseq X Plus with a 150 bp paired-end configuration. Adapter and quality trimming was performed using *bbduk* (sourceforge.net/projects/bbmap/). Reads were mapped using *bowtie2* (Langmead and Salzberg, 2012) on the GRCm38/mm10 genome, resulting sam files were converted to bam using *samtools* (Danecek et al., 2021), filtering and sorting was performed using *sambaba* (Tarasov et al., 2015) and indexing using *samtools*. Peak detection was performed using the *snakePipes* ATACseq pipeline and the default options (MACS2 as peak caller with qval < 0.001, CSAW for differential accessibility analysis with FDR < 0.05, logFC > 1). bigWig file of the three WT replicates and the three SoxC-KO replicates were merged into a single track using bigWigMerge for visualization in IGV (Robinson et al., 2011). Gene ontology analysis was performed using the enrichGO() function from the *clusterProfiler* v4.12.2 R package (Yu et al., 2012).

### Chromatin Immunoprecipitation DNA-sequencing

C57BL/6J E14.5 hindguts (minimum 40 hindguts per replicate, in 3 replicates) were dissected in PBS (gibco) as described above, divided into three replicates, digested 30 min at 37°C in 0.15 mg/mL collagenase D (Roche) and subsequently fixed in 1% paraformaldehyde (PFA, SantaCruz) for 10 minutes at room temperature. PFA was then quenched by adding glycine (Fluka) to a final concentration of 0.125 M. Cells were washed once with PBS, resuspended in 250 μL of lysis buffer (1% (SDS), 10 mM EDTA, 50 mM Tris-HCl pH 8, protease inhibitor cocktail (Roche)), frozen and thawed and incubated rotating for 30 min at 4°C. Lysates sonicated three minutes in Covaris S220 (intensity 5, duty cycle 10%, cycles per burst 200, frequency sweeping mode, temperature ca. 7°C) using AFA Fiber microTube (Covaris). Sonicated samples were diluted to 2.5 mL with dilution solution (0.01% SDS, 1% Triton X-100, 2 mM EDTA, 20 mM Tris-HCl pH 8, 150 mM NaCl, protease inhibitor cocktail (Roche)) and divided into 500 μL aliquotes. Fragmentation between 200 and 1000 bp was confirmed by running smaller aliquots on an agarose gel. Samples were incubated with antibodies (Table S2) overnight rotating at 4 °C, input samples were frozen for later de-crosslinking. Magna ChIP protein G magnetic beads (Sigma Aldrich) were blocked overnight with 0.05% BSA (Roche), 2 μg/ml of herring sperm DNA (Promega) in dilution solution. Antibody samples were then incubated with beads 3.5 h rotating at 4 °C and then washed with washing buffer (WB) 1 (0.1% SDS, 1% Tx100, 2 mM Tris-HCl pH 8, 150 mM NaCl), WB 2 (0.1% SDS, 1% Tx100, 2 mM Tris-HCl pH 8, 500 mM NaCl), WB 3 (1% NP40, 1% sodium deoxycholate, 10 mM Tris-HCl pH 8, 1 mM EDTA, 0.25 mM LiCl), and finally with WB 4 (10 mM Tris-HCl pH8 and 1 mM EDTA). WB 4 was removed and 100 μL of 10% CHELEX were added to the beads, 200 μL to the input samples and samples were de-crosslinked at 95°C for 10 min, cooled down at RT for 7 min, treated with Proteinase K (Fermentas, 2 μg/ml) at 55 °C for 30 min and inactivated at 95 °C for 10 min. Samples were centrifuged at max speed for 10 min, supernatant was collected. Beads and CHELEX pellet were washed with 100 μL water, centrifuged again and supernatant added to the previous one. For input samples, DNA precipitation was performed as follows: 28 μL of 5 M NaCl was added to 700 μL de-crosslinked input, 1400 μL 100% EtOH were added, and samples was incubated for at least 8 hours at -20 °C, centrifuged 30 min at 15’000 g, 4 °C, supernatant was removed, DNA pellet washed once with 70% EtOH and resuspended in a 100 μL water. Libraries were prepared by the FCGZ using the NEB Ultra II DNA Library Prep Kit (NEB) for Illumina according to the manufacturer’s instructions. For replicate 1, 200 M reads per sample were sequenced on an Illumina Novaseq 6000 with 150 bp paired end reads. For replicates 2 and 3, 20M reads per sample were sequenced on an Illumina Novaseq 6000 with 100 bp single read. Reads were mapped using the DNA-mapping pipeline from *snakePipes* pipelines, the premade GRCm38/mm10 Gencode release m19 indices and defaults options (BOWTIE2 as aligner). Peak detection was performed using the *snakePipes* ChIPseq pipeline and the default options (MACS2 as peak caller with qval<0.001). bigWig file of the three WT replicates and the three SoxC-KO replicates were merged into a single track using *bigWigMerge* for visualization in IGV. Peaks in at least two replicates out of three were found using the *bedops* suite (Neph et al., 2012) and the bedmap function with parameter --fraction-both 0.5 to return all genomic peak regions with at least 50% overlap in two files. Peaks were annotated using the R package *ChIPseeker* v1.40.0 (Yu et al., 2015) with the annotatePeak() function and the TxDb.Mmusculus.UCSC.mm10.knownGene database. Gene IDs in the Entrez format were translated to gene symbols using the mmusculus_gene_ensembl mart database and the getBM() R function of the *biomaRt* R package v2.60.1 (Durinck et al., 2005).

### Omics intersections

From differential analyses and ChIPseq annotated peak files, the unique gene symbols were extracted. For the oncoembryonic signature, genes downregulated in the adult compared to the embryo were intersected with genes upregulated in the cancer compared to the healthy. From ChIPseq identified peaks, unique gene symbols were extracted and the resulting gene list for Sox11 was merged with the one for Sox4. Annotations from regions with gained accessibility in ATACseq were filtered to obtain a gene symbol list with unique nearest gene entries, the same was done for the regions with lost accessibility. Gene symbols from transcripts down- or upregulated in SoxC-KO scRNAseq were extracted. All the resulting lists were used for the different intersections represented in Figure 2H. The Sankey plot was generated using SankeyMatic (https://sankeymatic.com/build/). For the SoxC oncoembryonic signature in Figure 3, genes downregulated in SoxC-KO and with an associated ATACseq region with lost accessibility in SoxC-KO were intersected with the oncoembryonic signature from Figure 1.

### Survival analyses

Kaplan-Meier plots for individual genes and the SoxC-oncoembryonic gene signature were generated using the Kaplan-Meier Plotter tool (https://kmplot.com/analysis/) (Györffy et al., 2010; Győrffy, 2024) with the colorectal cancer mRNA gene chip database. All patients were included, with parameters set to auto-select the best percentile cutoff and the JetSet best probe set (Li et al., 2011). Included datasets: GSE12945 (n=62), GSE13294 (n=155), GSE14333 (n=123), GSE143985 (n=91), GSE17538 (n=232), GSE18088 (n=53), GSE26682 (n=331), GSE30540 (n=35), GSE31595 (n=37), GSE33114 (n=90), GSE34489 (n=33), GSE37892 (n=65), GSE38832 (n=70), GSE39582 (n=514), GSE41285 (n=185), GSE92921 (n=59).

### Scoring of CRC patient’s scRNAseq data

scRNAseq data of CRC patients obtained from previously published datasets (GSE132465, GSE144735 (Lee et al., 2020) and GSE116222 (Parikh et al., 2019) only healthy subset). Standard pre-processing was performed as described above. Epithelial cells were subset based on published annotation. The signature score was calculated per cell using the AddModuleScore() function from the *Seurat* package. Adjusted scores were calculated by subtracting the mean score of all healthy cells. Averages per sample type (tumor, normal adjacent or healthy donor) and per patient were calculated and plotted using *ggplot2*.

### Total RNA extraction

Dissected tumors, healthy tissues or embryonic hindguts were resuspended in 300-500 μL of TRI Reagent (Sigma) and 60-100 μL of chloroform was added according to the manufacturer’s instruction. Samples were centrifuged for 15 min at 12’000 x g, and the aqueous phase was then mixed with one volume of 70% ethanol. The sample was loaded onto Qiagen RNeasy purification kit columns and purification continued as per manufacturer’s instruction including DNase treatment. 500 ng of total RNA was reverse-transcribed using the cDNA synthesis kit (Takara Bio) according to the manufacturer’s instructions. Expression of genes of interest was quantified with primers listed in Table S3, by RT–qPCR using the PowerTrack SYBR Green Master Mix (ThermoFisher) (2.5 ng of cDNA, 0.25 μM of each primer, 5 μL of master mix) and monitored by the QuantStudio3 system (Applied Biosystems). The samples were analyzed in technical duplicates and the average cycle threshold values were normalized to β-Actin using the ΔΔCT method (Livak and Schmittgen, 2001).

### Tissue isolation

Human patients’ samples, embryonic hindgut tissues from E14.5 embryos and colorectal tumors were fixed for 1 hour (overnight for RNAscope samples) at 4°C in 4% paraformaldehyde in PBS (ChemCruz) and incubated overnight in 30% sucrose in PBS at 4°C. Tissues were embedded in OCT and frozen at - 70°C.

### Cryo-sectioning

OCT-embedded tissues were cryo-sectioned using a CryoStar NX50 Cryostat (Thermo Scientific) at 10 μm and dried for 1 hour at room temperature before either being directly used for immunofluorescent staining or stored at -70°C.

### Immunofluorescent staining

Standard immunohistochemical protocols were performed with the primary antibodies listed in Table S2. Briefly, tissue sections were thawed at room temperature for 30 min and blocked with 5% BSA (Roche) + 0.2% Tween 20 (Sigma) in PBS (gibco). Primary antibodies were diluted in 3% BSA and incubated on the sections for 1 h at room temperature. After thorough washing with PBS, secondary antibodies (anti-rabbit, anti-goat antibodies conjugated with Alexa Fluor-555 and Alexa Fluor-647, Invitrogen) were diluted 1:500 dilution in 3% BSA and incubated on the sections for 1 h at room temperature. Phalloidin (Invitrogen, 1:1000) coupled to Alexa Fluor 488 was incubated together with secondary antibodies. Sections were counterstained with DAPI and, after washing with PBS, mounted with Dako Fluorescence Mounting Medium (Agilent) and imaged on the Leica SP8 confocal microscope maintained by the Center for Microscopy and Image Analysis, University of Zurich. Images were processed in FIJI (ImageJ 1.54f).

#### Image processing

The same adjustments were applied to all images to be compared. To quantify the nuclear fluorescent signal, a threshold (Huang algorithm, auto settings) was applied to the nuclear (DAPI) channel. The result was further binarized and improved by filling holes and applying a watershed filter. The nuclei were segmented using the analyze particle function (settings: 8-Infinity, circularity: 0–1). The segmentation was overlaid on the DAPI channel and all nuclei of low quality (e.g., partially sectioned/dim nuclei that were not fully captured) or of inconclusive position in regards to their tissue position were excluded. The average fluorescence intensity of the remaining nuclei in the proper channel was measured, plotted, and analyzed using Prism 8 (GraphPad, USA). To measure epithelium width, freehand lines were drawn in FIJI (ImageJ 1.54f) followed by length measurement.

For quantification of Klf4 and Sox11 signal along the spatial axis, a mask of the segmented nuclei was applied to the channel of interest, and all intensities outside of the nuclei were set to 0 using “Clear Outside”. A 1-pixel-wide region of interest along the x-axis was generated and the mean intensity of pixels with intensity > 1 across the region was measured. Intensities were plotted against the positional value in R using the barplot() function.

### RNA *in situ* hybridization

mRNA in situ hybridization was performed using the RNAscope technology™ (Advanced Cell Diagnostics, Germany) on mouse healthy adult colon and tumor tissue sections according to the manufacturer’s instructions (RNAscope Fluorescent Multiplex Assay). Probe sets for *Sox4* and *Sox11* were designed by Advanced Cell Diagnostics. Images were obtained on a Leica SP8 laser scanning confocal microscope.

### Luciferase assay

Tead2 enhancer region was cloned into pGL3 basic vector (AddGene 212936) using primers and enzymes listed in Table S4. Mutations in the binding site were induced by amplifying the full vector with mutation-containing primers. A minimal promoter was removed from the M50 Super 8x TOPFlash (AddGene 12456) by XhoI and HindIII digest and ligated to an XhoI digested pGL3. For the Sox11 overexpression plasmid, the Sox11 coding sequence was amplified from pLenti-CMV-GFP-Sox11 (AddGene, 120387). P2A-EGFP was amplified from a self-cloned sLP-CreERT2-P2A-EGFP plasmid. pLenti-Lifeact-EGFP (AddGene, 84383) was amplified using primers listed below. The three fragments were ligated using NEBuilder. Sox4-3myc- P2A sequence was PCR amplified from a synthesized plasmid and cloned into pLenti-Lifeact-tdTomato (AddGene, 64048) using NEBuilder. Klf4 was PCR amplified from the plasmid TetO-FUW-OSKM (AddGene, 20321) and inserted into XbaI-digested PB-CMV-MCS-EF1a-GreenPuro (SystemBiosciences, PB513B-1).

For luciferase measurements, 10K HEK- 293T cells were seeded in medium as described above in 96-well and after 10 h transfected with 17 ng enhancer/silencer pGL3 vector, 3.4 ng of pRL Renilla Luciferase vector (Promega, E2231) and 34 ng of transcription factor vector using the Lipofectamine™ 3000 reagent (Invitrogen) according to manufacturer’s instruction. Medium was replaced 12 h after transfection and 48 h after transfection cells were lysed with 50 μL 1x passive lysis buffer diluted in water (Promega, E194A). Cells were frozen at -70°C for 30 min. 10 μL of lysate was transferred to a white 96-well plate. 50 μL of Luciferase assay reagent II (Promega, E195A) and Stop&Glo reagent (Promega, E641A) were added to measure Firefly and Renilla successively in a GloMax-Multi microplate reader (Promega).

### Statistics

Images were prepared using Fiji (ImageJ, 1.52p), Adobe Illustrator (v28), Adobe Photoshop (v25) and Affinity Designer (v1.10). All statistical analyses were performed by Prism (GraphPad Software, v10, 2024). Statistical significances were measured using two-tailed unpaired t-test, one-way or two-way ANOVA with Tukey’s multiple comparison test. Relevant statistical information can be found in associated figure legends.

## Acknowledgments

We sincerely thank Konrad Basler for immense support including financing this work, George Hausmann for critical input on the manuscript, Erich Brunner, Luciano Rago, Sven Buchmann, Shyamala Maheswaran, Sakari Vanharanta, Ataman Sendoel, Darren Gilmour, Christoph Schneider and Claudio Cantù for treasured discussions, and members of the Basler lab for valuable input. We also thank Veronique Lefebvre and Elizabeth Sock for their generous gift of the SoxC conditional mouse model, Owen Sansom for kindly gifting the AKP organoids, Mark Robinson, Izaskun Mallona and Pierre-Luc Germain for bioinformatic counseling. We thank Susanna Salas for administrative assistance and Valerie Georgette Katimi Varela, Aline Böniger, Dillon King, Patrick Diener and Letizia Urietti for technical assistance. We also acknowledge the technical support from the Functional Genomics Center Zurich, especially Catharine Aquino, and from the Cytometry Facility of the University of Zurich, especially Mario Wickert and Manuel Schulthess. We also thank the Office for Animal Welfare and 3R of the University of Zurich, particularly Michaela Thallmair, Corina Berset, Nicole Wildner and Paulin Jirkof for supporting animal experimentation procedures. A.M. and R.J.P. are supported by the National Centres of Competence Molecular Systems Engineering (51NF40-182895). This work was supported by Swiss National Science Foundation (192475), University of Zurich Research Priority Program (URPP) “Translational Cancer Research”, Krebsliga (Swiss Cancer League) project funding (KFS-6031-02-2024). T.D. is supported by the Swiss 3R Competence Center (3RCC) (DP-2022-003) and by a grant from the Julius Klaus Stiftung. M.D.B. was supported by the Forschungskredit of the University of Zürich (FK-19-074). T.V. was supported by the project National Institute for Cancer Research (Programme EXCELES, ID Project No. LX22NPO5102)—Funded by the European Union— Next Generation EU. H.F. was supported by the University of Zurich GRC career grant (2024_Q1_CG_001), the Forschungskredit of the University of Zürich (FK-21-119) and a grant from the University and Medical Faculty of Zürich and the Comprehensive Cancer Center Zürich.

## Author contributions

Conceptualization: HF, Methodology: HF, TD, Investigation: HF, TD, GM, TV, MDB, AM, TB, SL, BMS, MN, Visualization: HF, TD, Supervision: HF, Resources: HF, MvdB, RJP, MS, Writing – original draft: HF, TD, Writing – review & editing: HF, TD, GM, TV, MDB, SL, MN, MvdB, AM, RJP, ICA

## Data and materials availability

Data and code are available upon request.

---

**Figure S1.**
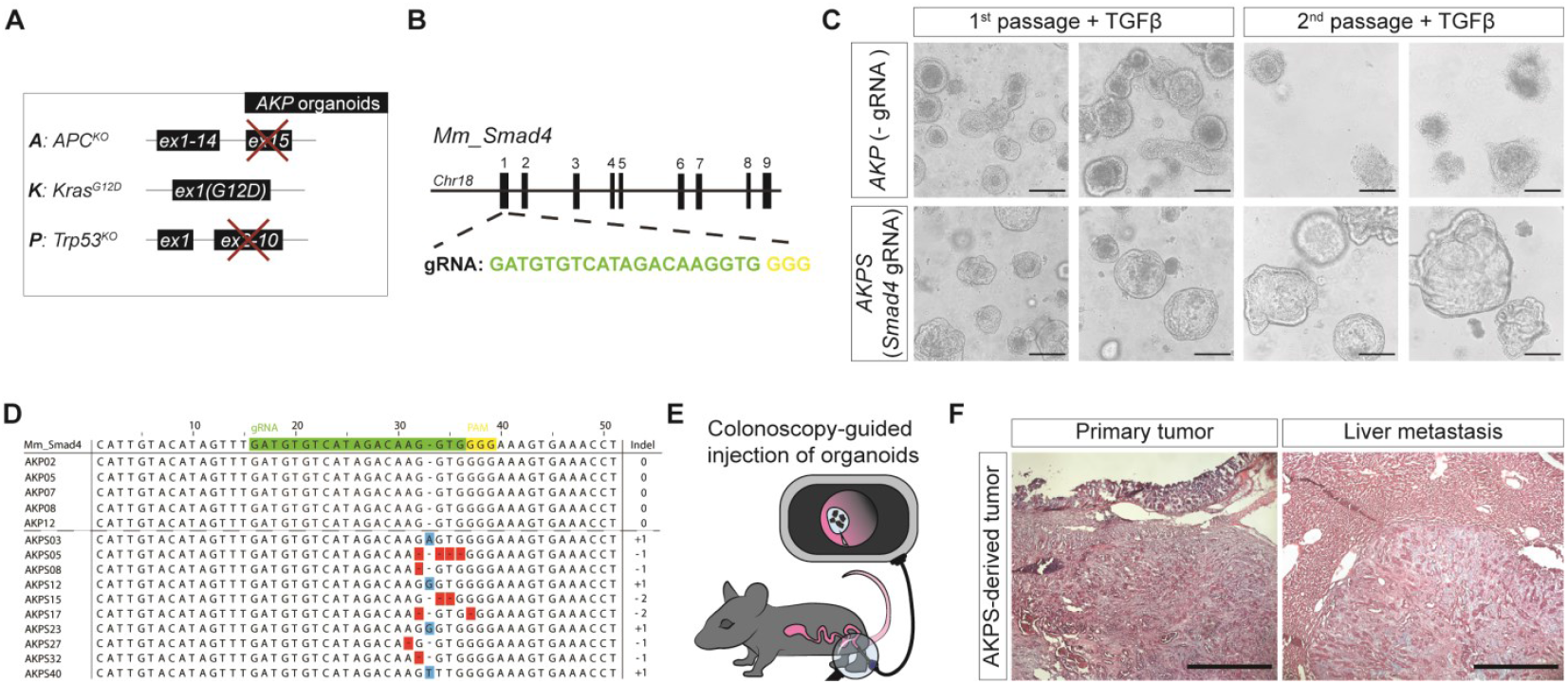
Engineering metastatic organoid-based CRC model by mutating *Smad4* in *Apc, Kras*, and *Tp53* organoid. (A) Schematic representation of the genetic background of AKP colorectal cancer (CRC) organoids, derived from a mouse model where Villin CreERT2-mediated recombination induces knockout of *Apc, Kras*, and *Tp53*. (B) CRISPR-Cas9 strategy used for *Smad4* mutagenesis, employing guide RNA (gRNA) targeting exon 1 of *Smad4*. (C) Comparison of control AKP organoids and AKP organoids with *Smad4* gRNA, selected with TGFβ recombinant protein. (D) Sanger sequencing results of clones generated from PCR amplifications of AKP and AKPS (AKP + *Smad4* mutation) organoids, confirming mutations in the *Smad4* gene. (E) Schematic of the orthotopic CRC model, where organoids are transplanted via colonoscopy-guided injection into the mouse colon. (F) Histological analysis of primary and metastatic tumors from the orthotopic CRC model visualized using hematoxylin and eosin staining.

**Figure S2.**
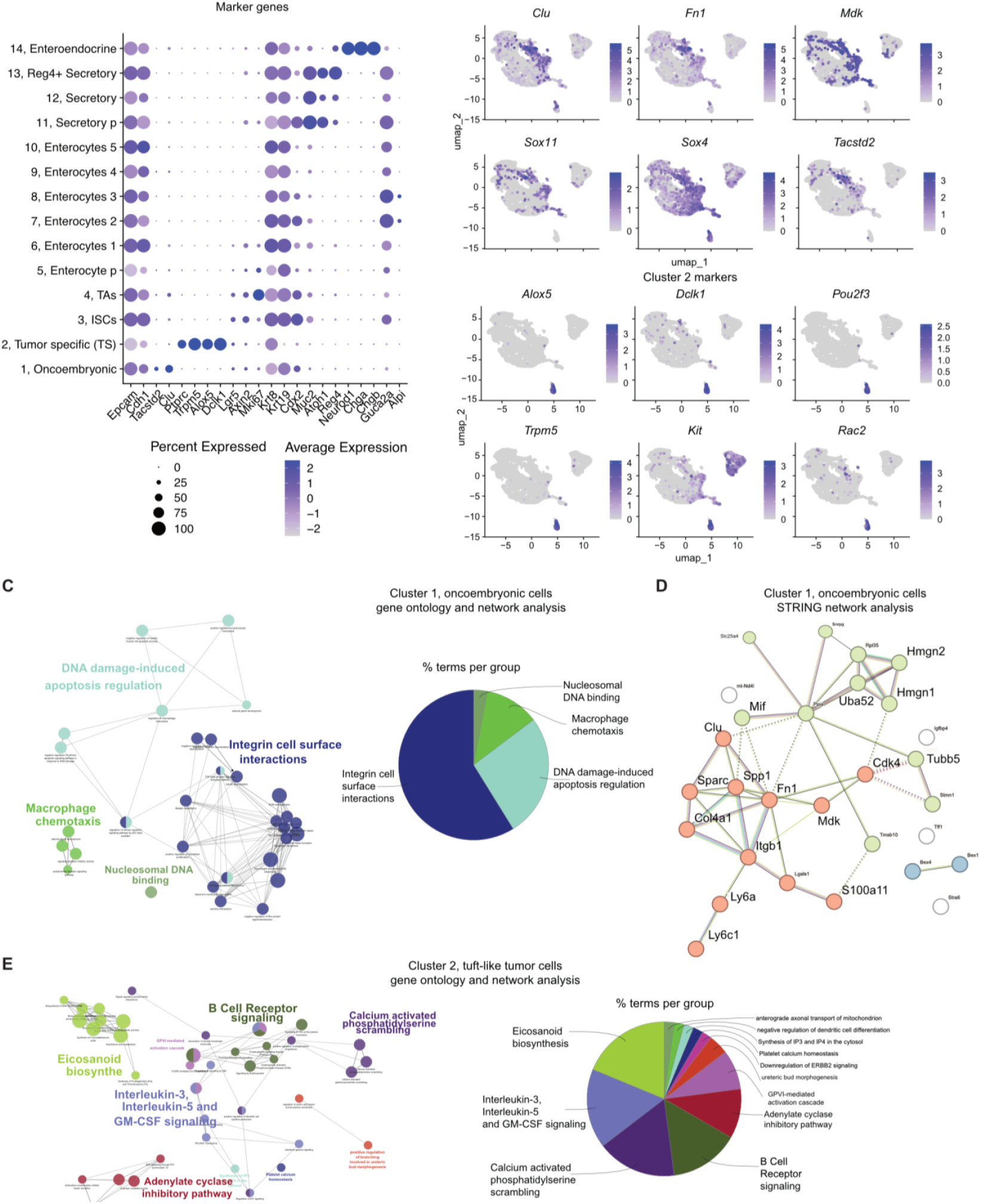
Tumor-specific cell populations exhibit distinct gene expression profiles and predicted functions. (A) Dot plot illustrating marker gene expression across annotated clusters from the single-cell RNA sequencing (scRNAseq) data presented in Figure 1A. The size of each dot represents the proportion of cells expressing a particular transcript, while the color intensity reflects the average expression level within each cluster. (B) UMAP plots from the scRNAseq data (see Figure1A) display the expression patterns of markers for cluster 1 and cluster 2. Color intensity, ranging from gray to purple, denotes the expression levels of each gene. (C) Predicted functional gene interaction network of highly expressed genes in cluster 1 (oncoembryonic cells) from the scRNAseq data in Figure 1A (left). A corresponding summary of gene ontology (GO) terms is shown as a pie chart (right), with numbers representing the percentage of GO terms for each category. (D) STRING functional protein association network analysis of highly expressed genes in cluster 1 cells from the scRNAseq data in Figure 1A. (E) Predicted functional gene interaction network of highly expressed genes in cluster 2 (tuft-like cells) from the scRNAseq data in Figure 1A (left), accompanied by a pie chart summarizing the GO terms (right), with numbers indicating the percentage of GO terms for each group.

**Figure S3.**
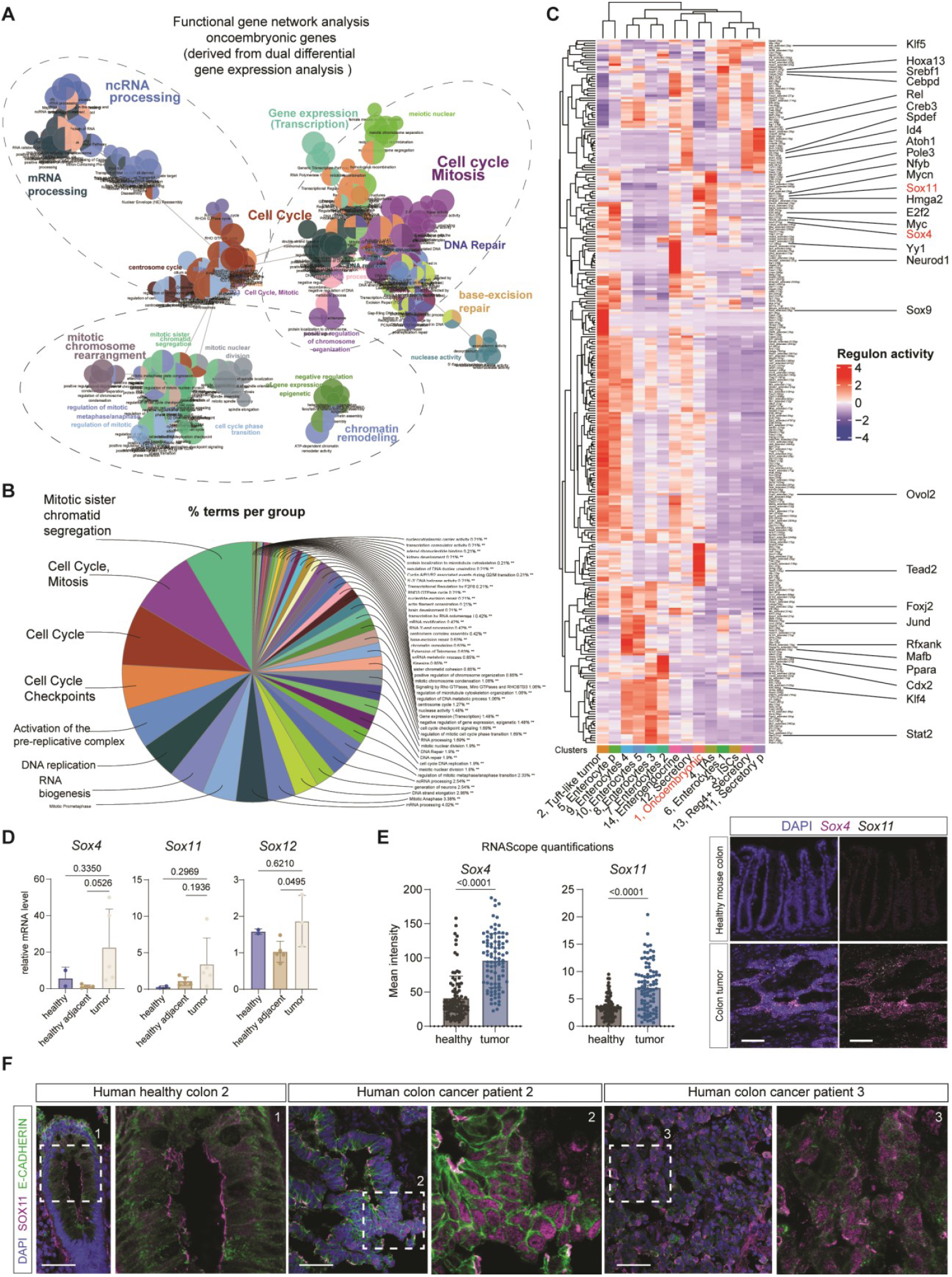
Sox4 and Sox11 as key drivers of transcription factor regulons enriched in oncoembryonic cells. (A) Predicted functional gene interaction network of oncoembryonic genes derived from dual differential gene expression analysis (from Figure 1B), showing significant interactions among key SoxC-regulated genes. (B) Summary of gene ontology (GO) terms represented in panel S3A, displayed as a pie chart, with numbers indicating the percentage of GO terms in each category. (C) Complete regulatory network inference using SCENIC analysis on integrated epithelial scRNAseq data (from Figure 1A), highlighting Sox4 and Sox11 as master regulators in the oncoembryonic cells. (D) Quantitative PCR showing transcript levels of SoxC transcription factor targets in healthy, healthy-adjacent, and colon tumor tissues from mouse models. (E) RNA in situ hybridization (RNAscope) and quantification of signal intensity for Sox4 and Sox11 in mouse healthy and colon tumor tissues. Scale bar: 50 μm, DAPI (blue) counterstains nuclei. (F) Immunofluorescent staining of human healthy and cancerous colon tissues for E-CADHERIN (epithelial marker) and SOX11 protein. Scale bar: 50 μm.

**Figure S4.**
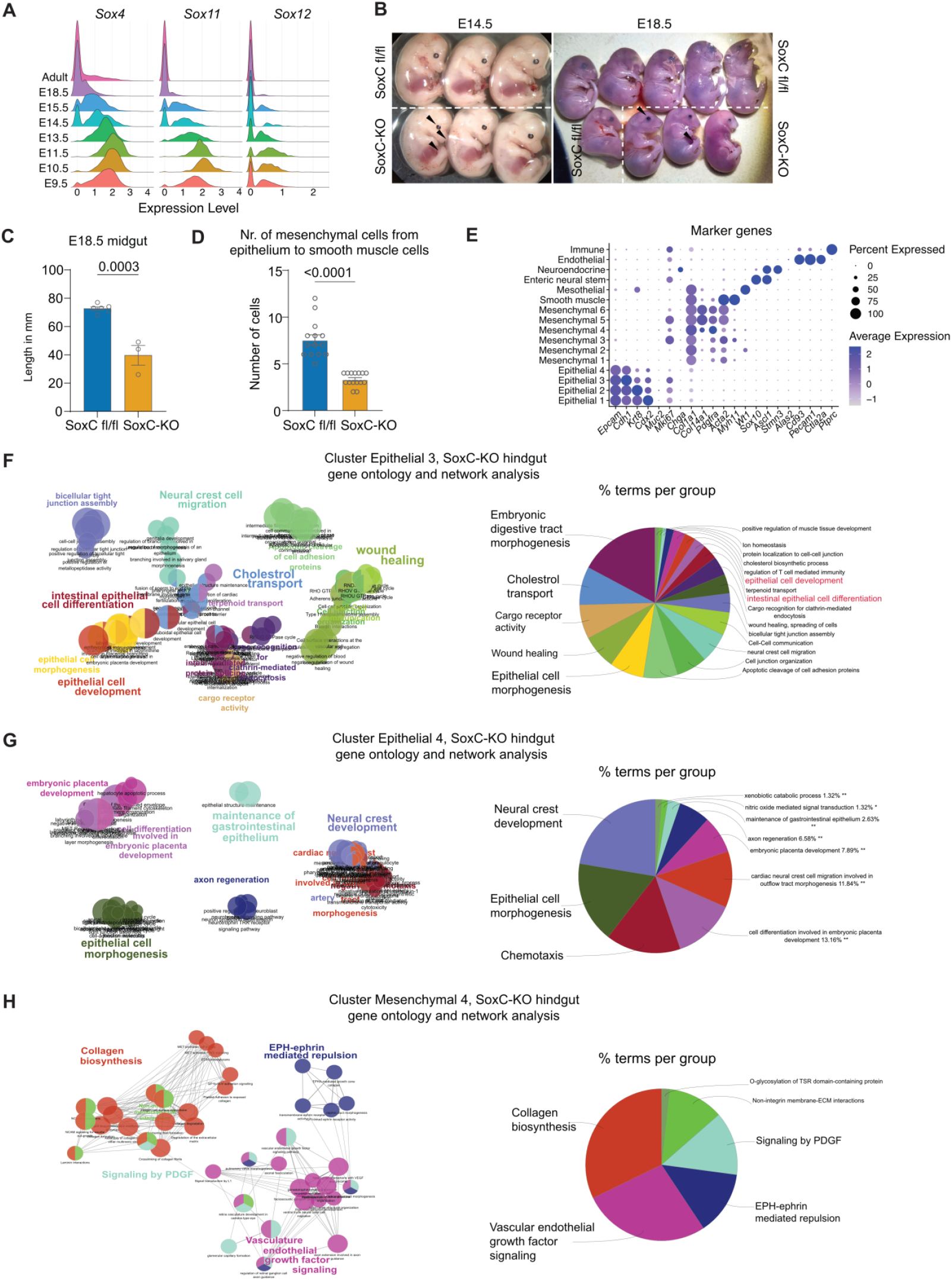
SoxC transcription factors control colon development. (A) Ridge plot representing normalized counts of *Sox11, Sox4* and *Sox12* in epithelial cells at different stages of colon development. (B) Full R26-CreERT2; SoxC fl/fl or non-Cre littermate embryos at stage E14.5 or E18.5 after tamoxifen induction of Cre at E10.5. Arrows point at eye, back and digit phenotypes. (C) Length (in mm) of SoxC-KO or non-Cre littermate midguts at E18.5. (D) Quantification 29 of mesenchymal nuclei present in the region below the epithelium and before the smooth muscle layer in SoxC-KO or non-Cre littermate E14.5 hindguts. (E) Marker genes were used for the annotation of the integrated WT and SoxC-KO E14.5 hindgut scRNAseq dataset. (F-H) Gene ontology and network analyses of top 200 genes enriched in clusters Epithelial 3 (F), Epithelial 4 (G) and Mesenchymal 4 (H) of the integrated WT and SoxC-KO E14.5 hindgut scRNAseq dataset.

**Figure S5.**
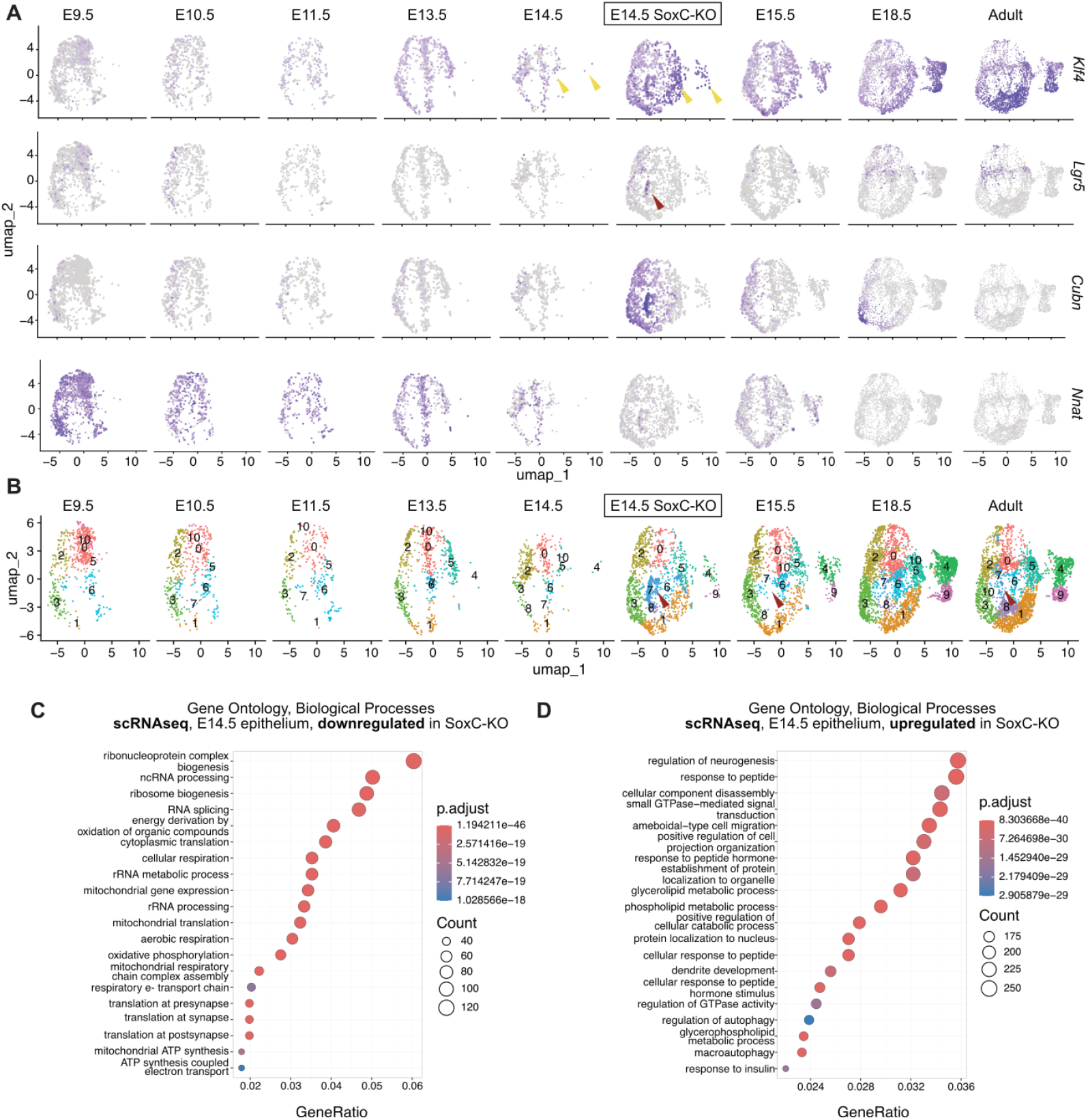
Premature differentiation and other biological processes induced by SoxC-KO. (A) UMAP plot of epithelial cells at different timepoints of hindgut development showing expression of selected genes. Yellow arrows point at adult-like cells prematurely appearing in SoxC-KO E14.5 hindguts. Red arrow points at cell cluster enriched in SoxC- KO E14.5 hindgut. (B) UMAP plot of epithelial cells at different time points of hindgut development. Single cells are colored based on cluster annotation. Red arrow points at the cell cluster enriched in SoxC-KO E14.5 hindgut as in (B). (C-D) Gene ontology analysis (biological processes) of all genes downregulated (C) or upregulated (D) in SoxC-KO E14.5 hindgut compared to WT. The top 20 gene ontologies are shown.

**Figure S6.**
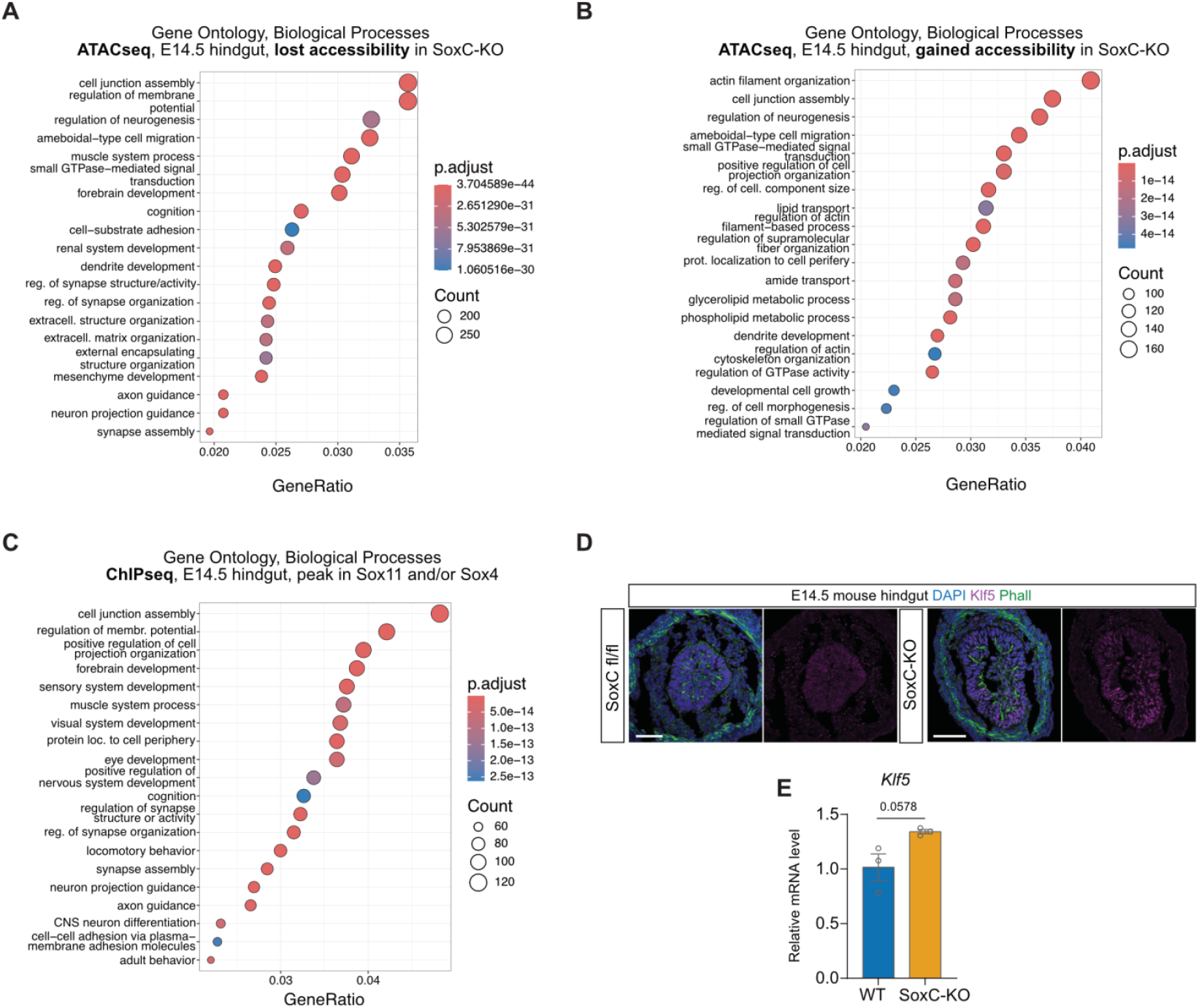
Biological processes affected by SoxC-KO in ATACseq and ChIPseq. (A-C) Gene ontology analysis (biological processes) of all genes nearest to genomic regions with lost accessibility (A) or gained accessibility (B) in SoxC-KO E14.5 hindgut compared to WT or with significant ChIPseq peaks for Sox11 and or Sox4 (C). Top 20 gene ontologies are shown. (D) Representative immunofluorescent staining in SoxC-KO or non-Cre littermate E14.5 hindguts. Blue DAPI, magenta Klf5, green Phalloidin. Scalebar 50 μm. (E) qPCR analysis of *Klf5* expression in WT and SoxC-KO E14.5 hindguts. Statistical analysis was performed using a two-tailed unpaired t-test.

**Figure S7.**
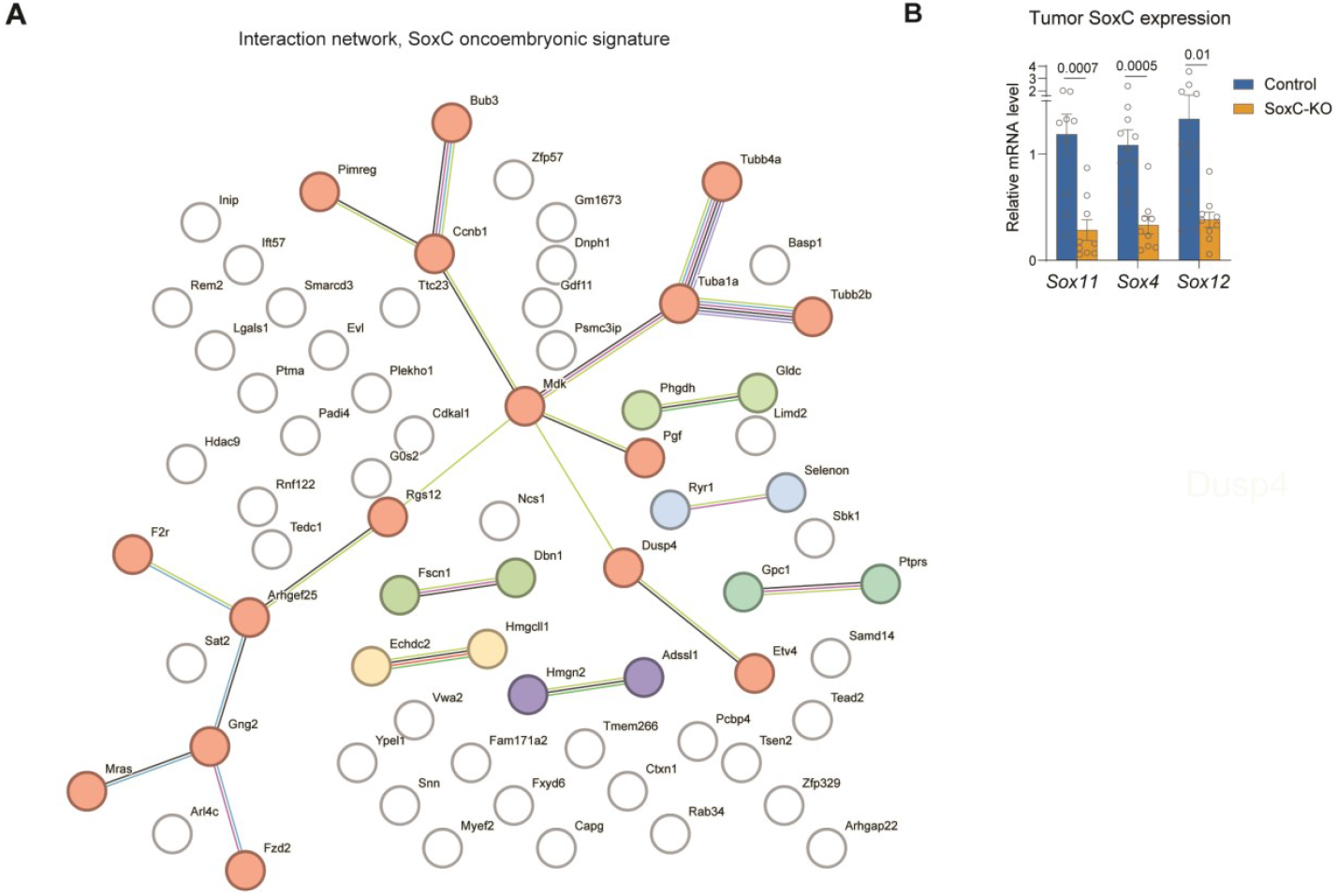
Functional protein interaction network of the SoxC oncoembryonic signature. **(A)** STRING functional protein association network analysis of the 71-gene SoxC oncoembryonic program signature (from Figure 3A), illustrating predicted interactions between key gene products involved in the SoxC-driven pathway. **(B)** Quantitative PCR analysis displaying transcript levels of selected SoxC transcription factor targets in control and SoxC-KO tumor samples. The results highlight the impact of SoxC deletion on downstream gene expression.

**Figure S8.**
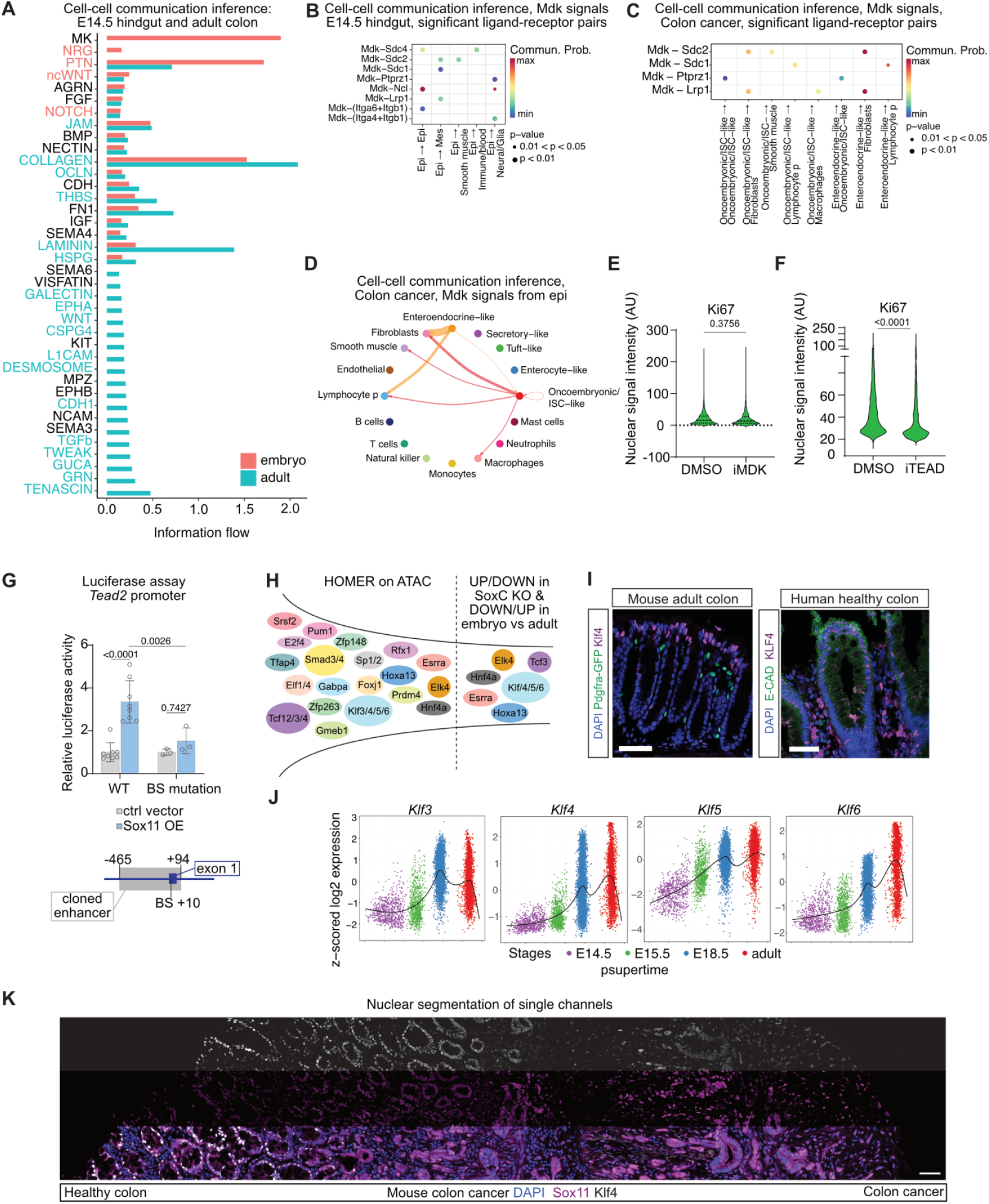
*Mdk* and *Tead2* as key downstream targets of SoxC TFs. (A) Comparison of inferred information flow for all predicted cell-communication pathways in E14.5 hindgut or adult colon. Red-colored pathways are enriched in the E14.5 hindgut, blue-colored pathways are enriched in the adult. (B) Significant predicted Mdk ligand-receptor interactions from cell-cell communication prediction by CellChat in the scRNAseq of E14.5 WT including all cell types. Edge weights are proportional to the interaction strength. (C) Significant predicted Mdk ligand-receptor interactions originating from epithelial cells (Oncoembryonic/ISC-like, Enterocyte-like, Tuft-like, Secretory-like or Enteroendocrine- Dalessi et al., 2024 (preprint) like). Inferred interactions are the result of cell-cell communication prediction by CellChat in the scRNAseq of cancer cells only (AKPS organoid-derived tumors) including all cell types. Edge weights are proportional to the interaction strength. (D) Inferred interactions through Mdk signaling from epithelial cells as in (B). (E) Quantification of nuclear Ki67 signal in Figure 5D and replicates (E13.5 hindgut explants cultured for 2 days *ex vivo* in the presence of DMSO or MDK inhibitor). (F) Quantification of nuclear Ki67 signal in Figure 5G and replicates (E12.5 hindgut explants cultured for 2 days *ex vivo* in the presence of DMSO or TEAD inhibitor). (G) Top: Activity of WT *Tead2* promoter or with deletion of putative Sox11 binding site (BS) after Sox11 transfection assessed by luciferase assay in HEK293 cells. N=8 for WT, n=3 for BS mutation. Bottom: schematics of *Tead2* promoter region cloned for the assay (−465 to +94 from the coding sequence start) and the mutated base (position +10 from the coding sequence start). Statistical analysis using two-way Anova with Bonferroni’s multiple comparison test. (H) Schematics of strategy to identify intermediate regulators of indirect targets of SoxC TFs. (I) Representative immunofluorescent staining of adult mouse colon (left, blue DAPI, green Pdgfra-GFP, magenta Klf4) or human healthy colon (right, blue DAPI, green E-CADHERIN, magenta KLF4). (J) Z- scored log2 expression values of *Klf3, Klf4, Klf5, Klf6* with cell ordered along a pseudotime learned by *pstupertime* at different stages of colon development (x-axis). (K) Immunofluorescent staining from Figure 5M (mouse colon cancer) with nuclear mask applied on each separate channel. Top: grey, Klf4. Middle: magenta, Sox11. Bottom: blue DAPI, grey Klf4, magenta Sox11. If not otherwise stated, statistical analyses were performed using a two-tailed unpaired t-test. Scale bars 50 μm.

## References

Agbo, K.C., Huang, J.Z., Ghaleb, A.M., Williams, J.L., Shroyer, K.R., Bialkowska, A.B., Yang, V.W., 2019. Loss of the Krüppel-like factor 4 tumor suppressor is associated with epithelial-mesenchymal transition in colorectal cancer. jcmt 5, N/A-N/A. 10.20517/2394-4722.2019.35

Aibar, S., González-Blas, C.B., Moerman, T., Huynh-Thu, V.A., Imrichova, H., Hulselmans, G., Rambow, F., Marine, J.-C., Geurts, P., Aerts, J., van den Oord, J., Atak, Z.K., Wouters, J., Aerts, S., 2017. SCENIC: single-cell regulatory network inference and clustering. Nat Methods 14, 1083–1086. 10.1038/nmeth.4463

Aiello, N.M., Stanger, B.Z., 2016. Echoes of the embryo: using the developmental biology toolkit to study cancer. Disease Models & Mechanisms 9, 105– 114. 10.1242/dmm.023184

AR Moorman, F Cambuli, EK Benitez, Q Jiang, Y Xie, A Mahmoud, M Lumish, S Hartner, S Balkaran, J Bermeo, S Asawa, C Firat, A Saxena, A Luthra, V Sgambati, K Luckett, F Wu, Y Li, Z Yi, I Masilionis, K Soares, E Pappou, R Yaeger, P Kingham, W Jarnagin, P Paty, MR Weiser, L Mazutis, M D’Angelica, J Shia, J Garcia-Aguilar, T Nawy, TJ Hollmann, R Chaligné, F Sanchez-Vega, R Sharma, D Pe’er, K Ganesh, 2023. Progressive plasticity during colorectal cancer metastasis. bioRxiv 2023.08.18.553925. 10.1101/2023.08.18.553925

Bala, P., Rennhack, J.P., Aitymbayev, D., Morris, C., Moyer, S.M., Duronio, G.N., Doan, P., Li, Z., Liang, X., Hornick, J.L., Yurgelun, M.B., Hahn, W.C., Sethi, N.S., 2023. Aberrant cell state plasticity mediated by developmental reprogramming precedes colorectal cancer initiation. Science Advances 9, eadf0927. 10.1126/sciadv.adf0927

Baulies, A., Moncho-Amor, V., Drago-Garcia, D., Kucharska, A., Chakravarty, P., Moreno-Valladares, M., Cruces-Salguero, S., Hubl, F., Hutton, C., Kim, H., Matheu, A., Lovell-Badge, R., Li, V.S.W., 2024. SOX2 drives fetal reprogramming and reversible dormancy in colorectal cancer. bioRxiv 2024.09.11.612412. 10.1101/2024.09.11.612412

Bergsland, M., Werme, M., Malewicz, M., Perlmann, T., Muhr, J., 2006. The establishment of neuronal properties is controlled by Sox4 and Sox11. Genes Dev 20, 3475–3486. 10.1101/gad.403406

Bhattaram, P., Penzo-Méndez, A., Sock, E., Colmenares, C., Kaneko, K.J., Vassilev, A., DePamphilis, M.L., Wegner, M., Lefebvre, V., 2010. Organogenesis relies on SoxC transcription factors for the survival of neural and mesenchymal progenitors. Nature Communications 1, 1–12. 10.1038/ncomms1008

Biller, L.H., Schrag, D., 2021. Diagnosis and Treatment of Metastatic Colorectal Cancer: A Review. JAMA 325, 669–685. 10.1001/jama.2021.0106

Bindea, G., Mlecnik, B., Hackl, H., Charoentong, P., Tosolini, M., Kirilovsky, A., Fridman, W.-H., Pagès, F., Trajanoski, Z., Galon, J., 2009. ClueGO: a Cytoscape plug-in to decipher functionally grouped gene ontology and pathway annotation networks. Bioinformatics 25, 1091–1093. 10.1093/bioinformatics/btp101

Borrelli, C., Roberts, M., Eletto, D., Hussherr, M.-D., Fazilaty, H., Valenta, T., Lafzi, A., Kretz, J.A., Guido Vinzoni, E., Karakatsani, A., Adivarahan, S., Mannhart, A., Kimura, S., Meijs, A., Baccouche Mhamedi, F., Acar, I.E., Handler, K., Ficht, X., Platt, R.J., Piscuoglio, S., Moor, A.E., 2024. In vivo interaction screening reveals liver-derived constraints to metastasis. Nature 632, 411–418. 10.1038/s41586-024-07715-3

Brügger, M.D., Tomas, V., Fazilaty, H., George, H., Konrad, B., 2020. Distinct populations of crypt-associated fibroblasts act as signaling hubs to control colon homeostasis. PloS Biol.

Buenrostro, J.D., Wu, B., Chang, H.Y., Greenleaf, W.J., 2015. ATAC-seq: A Method for Assaying Chromatin Accessibility Genome-Wide. Current Protocols in Molecular Biology 109, 21.29.1-21.29.9. 10.1002/0471142727.mb2129s109

Burdziak, C., Alonso-Curbelo, D., Walle, T., Reyes, J., Barriga, F.M., Haviv, D., Xie, Y., Zhao, Z., Zhao, C.J., Chen, H.-A., Chaudhary, O., Masilionis, I., Choo, Z.-N., Gao, V., Luan, W., Wuest, A., Ho, Y.-J., Wei, Y., Quail, D.F., Koche, R., Mazutis, L., Chaligné, R., Nawy, T., Lowe, S.W., Pe’er, D., 2023. Epigenetic plasticity cooperates with cell-cell interactions to direct pancreatic tumorigenesis. Science 380, eadd5327. 10.1126/science.add5327

Butler, A., Hoffman, P., Smibert, P., Papalexi, E., Satija, R., 2018. Integrating single-cell transcriptomic data across different conditions, technologies, and species. Nat. Biotechnol. 36, 411–420. 10.1038/nbt.4096

Campa, C.C., Weisbach, N.R., Santinha, A.J., Incarnato, D., Platt, R.J., 2019. Multiplexed genome engineering by Cas12a and CRISPR arrays encoded on single transcripts. Nat Methods 16, 887–893. 10.1038/s41592-019-0508-6

Cano, A., Pérez-Moreno, M.A., Rodrigo, I., Locascio, A., Blanco, M.J., del Barrio, M.G., Portillo, F., Nieto, M.A., 2000. The transcription factor Snail controls epithelial–mesenchymal transitions by repressing E-cadherin expression. Nat Cell Biol 2, 76–83. 10.1038/35000025

Cerezo-Wallis, D., Contreras-Alcalde, M., Troulé, K., Catena, X., Mucientes, C., Calvo, T.G., Cañón, E., Tejedo, C., Pennacchi, P.C., Hogan, S., Kölblinger, P., Tejero, H., Chen, A.X., Ibarz, N., Graña-Castro, O., Martinez, L., Muñoz, J., Ortiz-Romero, P., Rodriguez-Peralto, J.L., Gómez-López, G., Al-Shahrour, F., Rabadán, R., Levesque, M.P., Olmeda, D., Soengas, M.S., 2020. Midkine rewires the melanoma microenvironment toward a tolerogenic and immune-resistant state. Nat Med 26, 1865–1877. 10.1038/s41591-020-1073-3

Danecek, P., Bonfield, J.K., Liddle, J., Marshall, J., Ohan, V., Pollard, M.O., Whitwham, A., Keane, T., McCarthy, S.A., Davies, R.M., Li, H., 2021. Twelve years of SAMtools and BCFtools. GigaScience 10, giab008. 10.1093/gigascience/giab008

de Sousa e Melo, F., Kurtova, A.V., Harnoss, J.M., Kljavin, N., Hoeck, J.D., Hung, J., Anderson, J.E., Storm, E.E., Modrusan, Z., Koeppen, H., Dijkgraaf, G.J.P., Piskol, R., de Sauvage, F.J., 2017. A distinct role for Lgr5+ stem cells in primary and metastatic colon cancer. Nature 543, 676–680. 10.1038/nature21713

DeWeirdt, P.C., Sanson, K.R., Sangree, A.K., Hegde, M., Hanna, R.E., Feeley, M.N., Griffith, A.L., Teng, T., Borys, S.M., Strand, C., Joung, J.K., Kleinstiver, B.P., Pan, X., Huang, A., Doench, J.G., 2021. Optimization of AsCas12a for combinatorial genetic screens in human cells. Nat Biotechnol 39, 94–104. 10.1038/s41587-020-0600-6

Dobin, A., Davis, C.A., Schlesinger, F., Drenkow, J., Zaleski, C., Jha, S., Batut, P., Chaisson, M., Gingeras, T.R., 2013. STAR: ultrafast universal RNA-seq aligner. Bioinformatics 29, 15–21. 10.1093/bioinformatics/bts635

Durinck, S., Moreau, Y., Kasprzyk, A., Davis, S., De Moor, B., Brazma, A., Huber, W., 2005. BioMart and Bioconductor: a powerful link between biological databases and microarray data analysis. Bioinformatics 21, 3439–3440. 10.1093/bioinformatics/bti525

Fang, T., Liang, T., Wang, Y., Wu, H., Liu, S., Xie, L., Liang, J., Wang, C., Tan, Y., 2021. Prognostic role and clinicopathological features of SMAD4 gene mutation in colorectal cancer: a systematic review and meta-analysis. BMC Gastroenterol 21, 297. 10.1186/s12876-021-01864-9

Fazilaty, H., 2023. Restoration of embryonic gene expression patterns in tissue regeneration and disease. Nat Rev Mol Cell Biol 1–2. 10.1038/s41580-023-00586-y

Fazilaty, H., Basler, K., 2023. Reactivation of embryonic genetic programs in tissue regeneration and disease. Nat Genet 55, 1792–1806. 10.1038/s41588-023-01526-4

Fazilaty, H., Brügger, M.D., Valenta, T., Szczerba, B.M., Berkova, L., Doumpas, N., Hausmann, G., Scharl, M., Basler, K., 2021. Tracing colonic embryonic transcriptional profiles and their reactivation upon intestinal damage. Cell Reports 36. 10.1016/j.celrep.2021.109484

Fazilaty, H., Rago, L., Kass Youssef, K., Ocaña, O.H., Garcia-Asencio, F., Arcas, A., Galceran, J., Nieto, M.A., 2019. A gene regulatory network to control EMT programs in development and disease. Nature Communications 10, 1–16. 10.1038/s41467-019-13091-8

Fu, L., Shi, Y.-B., 2017. The Sox transcriptional factors: Functions during intestinal development in vertebrates. Seminars in Cell & Developmental Biology, Regulation of development by SOX proteins 63, 58–67. 10.1016/j.semcdb.2016.08.022

Gold, P., Freedman, S.O., 1965. SPECIFIC CARCINOEMBRYONIC ANTIGENS OF THE HUMAN DIGESTIVE SYSTEM. Journal of Experimental Medicine 122, 467–481. 10.1084/jem.122.3.467

Győrffy, B., 2024. Integrated analysis of public datasets for the discovery and validation of survival-associated genes in solid tumors. Innovation 5. 10.1016/j.xinn.2024.100625

Györffy, B., Lanczky, A., Eklund, A.C., Denkert, C., Budczies, J., Li, Q., Szallasi, Z., 2010. An online survival analysis tool to rapidly assess the effect of 22,277 genes on breast cancer prognosis using microarray data of 1,809 patients. Breast Cancer Res Treat 123, 725–731. 10.1007/s10549-009-0674-9

Hanahan, D., 2022. Hallmarks of Cancer: New Dimensions. Cancer Discovery 12, 31–46. 10.1158/2159-8290.CD-21-1059

Hanahan, D., Weinberg, R.A., 2011. Hallmarks of Cancer: The Next Generation. Cell 144, 646–674. 10.1016/j.cell.2011.02.013

Hashimoto, M., Kojima, Y., Sakamoto, T., Ozato, Y., Nakano, Y., Abe, T., Hosoda, K., Saito, H., Higuchi, S., Hisamatsu, Y., Toshima, T., Yonemura, Y., Masuda, T., Hata, T., Nagayama, S., Kagawa, K., Goto, Y., Utou, M., Gamachi, A., Imamura, K., Kuze, Y., Zenkoh, J., Suzuki, A., Takahashi, K., Niida, A., Hirose, H., Hayashi, S., Koseki, J., Fukuchi, S., Murakami, K., Yoshizumi, T., Kadomatsu, K., Tobo, T., Oda, Y., Uemura, M., Eguchi, H., Doki, Y., Mori, M., Oshima, M., Shibata, T., Suzuki, Y., Shimamura, T., Mimori, K., 2024. Spatial and single-cell colocalisation analysis reveals MDK-mediated immunosuppressive environment with regulatory T cells in colorectal carcinogenesis. eBioMedicine 103. 10.1016/j.ebiom.2024.105102

Hickey, J.W., Becker, W.R., Nevins, S.A., Horning, A., Perez, A.E., Zhu, C., Zhu, B., Wei, B., Chiu, R., Chen, D.C., Cotter, D.L., Esplin, E.D., Weimer, A.K., Caraccio, C., Venkataraaman, V., Schürch, C.M., Black, S., Brbić, M., Cao, K., Chen, S., Zhang, W., Monte, E., Zhang, N.R., Ma, Z., Leskovec, J., Zhang, Z., Lin, S., Longacre, T., Plevritis, S.K., Lin, Y., Nolan, G.P., Greenleaf, W.J., Snyder, M., 2023. Organization of the human intestine at single-cell resolution. Nature 619, 572–584. 10.1038/s41586-023-05915-x

Hoser, M., Potzner, M.R., Koch, J.M.C., Bösl, M.R., Wegner, M., Sock, E., 2008. Sox12 Deletion in the Mouse Reveals Nonreciprocal Redundancy with the Related Sox4 and Sox11 Transcription Factors. Molecular and Cellular Biology 28, 4675–4687. 10.1128/MCB.00338-08

Hua, Q., Sun, Z., Liu, Y., Shen, X., Zhao, W., Zhu, X., Xu, P., 2021. KLK8 promotes the proliferation and metastasis of colorectal cancer via the activation of EMT associated with PAR1. Cell Death Dis 12, 1–14. 10.1038/s41419-021-04149-x

Huynh-Thu, V.A., Irrthum, A., Wehenkel, L., Geurts, P., 2010. Inferring Regulatory Networks from Expression Data Using Tree-Based Methods. PLOS ONE 5, e12776. 10.1371/journal.pone.0012776

Jin, S., Guerrero-Juarez, C.F., Zhang, L., Chang, I., Ramos, R., Kuan, C.-H., Myung, P., Plikus, M.V., Nie, Q., 2021. Inference and analysis of cell-cell communication using CellChat. Nat Commun 12, 1088. 10.1038/s41467-021-21246-9

Jin, S., Plikus, M.V., Nie, Q., 2023. CellChat for systematic analysis of cell-cell communication from single-cell and spatially resolved transcriptomics. 10.1101/2023.11.05.565674

Kaminow, B., Yunusov, D., Dobin, A., 2021. STARsolo: accurate, fast and versatile mapping/quantification of single-cell and single-nucleus RNA-seq data. bioRxiv 2021.05.05.442755. 10.1101/2021.05.05.442755

Katz, J.P., Perreault, N., Goldstein, B.G., Lee, C.S., Labosky, P.A., Yang, V.W., Kaestner, K.H., 2002. The zinc-finger transcription factor Klf4 is required for terminal differentiation of goblet cells in the colon. Development 129, 2619–2628. 10.1242/dev.129.11.2619

Kim, H.K., Min, S., Song, M., Jung, S., Choi, J.W., Kim, Y., Lee, S., Yoon, S., Kim, H. (Henry), 2018. Deep learning improves prediction of CRISPR–Cpf1 guide RNA activity. Nat Biotechnol 36, 239–241. 10.1038/nbt.4061

Kleinstiver, B.P., Sousa, A.A., Walton, R.T., Tak, Y.E., Hsu, J.Y., Clement, K., Welch, M.M., Horng, J.E., Malagon-Lopez, J., Scarfò, I., Maus, M.V., Pinello, L., Aryee, M.J., Joung, J.K., 2019. Engineered CRISPR-Cas12a variants with increased activities and improved targeting ranges for gene, epigenetic and base editing. Nat Biotechnol 37, 276–282. 10.1038/s41587-018-0011-0

Krebs, E.T., 1947. CANCER AND THE EMBRYONAL HYPOTHESIS. Calif Med 66, 270–271.

Langmead, B., Salzberg, S.L., 2012. Fast gapped-read alignment with Bowtie 2. Nat Methods 9, 357– 359. 10.1038/nmeth.1923

Lee, H.-O., Hong, Y., Etlioglu, H.E., Cho, Y.B., Pomella, V., Van den Bosch, B., Vanhecke, J., Verbandt, S., Hong, H., Min, J.-W., Kim, N., Eum, H.H., Qian, J., Boeckx, B., Lambrechts, D., Tsantoulis, P., De Hertogh, G., Chung, W., Lee, T., An, M., Shin, H.-T., Joung, J.-G., Jung, M.-H., Ko, G., Wirapati, P., Kim, S.H., Kim, H.C., Yun, S.H., Tan, I.B.H., Ranjan, B., Lee, W.Y., Kim, T.-Y., Choi, J.K., Kim, Y.-J., Prabhakar, S., Tejpar, S., Park, W.-Y., 2020. Lineage-dependent gene expression programs influence the immune landscape of colorectal cancer. Nature Genetics 52, 594–603. 10.1038/s41588-020-0636-z

Li, Q., Birkbak, N.J., Gyorffy, B., Szallasi, Z., Eklund, A.C., 2011. Jetset: selecting the optimal microarray probe set to represent a gene. BMC Bioinformatics 12, 474. 10.1186/1471-2105-12-474

Liu, Y.-N., Abou-Kheir, W., Yin, J.J., Fang, L., Hynes, P., Casey, O., Hu, D., Wan, Y., Seng, V., Sheppard-Tillman, H., Martin, P., Kelly, K., 2012. Critical and Reciprocal Regulation of KLF4 and SLUG in Transforming Growth Factor β-Initiated Prostate Cancer Epithelial-Mesenchymal Transition. Molecular and Cellular Biology 32, 941–953. 10.1128/MCB.06306-11

Livak, K.J., Schmittgen, T.D., 2001. Analysis of relative gene expression data using real-time quantitative PCR and the 2(-Delta Delta C(T)) Method. Methods 25, 402–408. 10.1006/meth.2001.1262

Macnair, W., Gupta, R., Claassen, M., 2022. psupertime: supervised pseudotime analysis for time-series single-cell RNA-seq data. Bioinformatics 38, i290–i298. 10.1093/bioinformatics/btac227

Mallona, I., Robinson, M.D., 2024. Method code for RoCK and ROI: a single-cell RNA-sequencing method with enhanced transcriptome information via targeted enrichment of transcripts and preselected, region-specific sequencing. 10.5281/zenodo.11070201

McInnes, L., Healy, J., Melville, J., 2018. UMAP: Uniform Manifold Approximation and Projection for Dimension Reduction. 1802.03426 [cs, stat].

Miao, Q., Hill, M.C., Chen, F., Mo, Q., Ku, A.T., Ramos, C., Sock, E., Lefebvre, V., Nguyen, H., 2019. SOX11 and SOX4 drive the reactivation of an embryonic gene program during murine wound repair. Nature Communications 10, 4042. 10.1038/s41467-019-11880-9

Moro, G., Mallona, I., Maillard, J., Brügger, M.D., Fazilaty, H., Szabo, Q., Valenta, T., Handler, K., Kerlin, F., Moor, A.E., Zinzen, R., Robinson, M.D., Brunner, E., Basler, K., 2024. RoCK and ROI: Single-cell transcriptomics with multiplexed enrichment of selected transcripts and region-specific sequencing. 10.1101/2024.05.18.594120

Mustata, R.C., Vasile, G., Fernandez-Vallone, V., Strollo, S., Lefort, A., Libert, F., Monteyne, D., Pérez-Morga, D., Vassart, G., Garcia, M.-I., 2013. Identification of Lgr5-independent spheroid-generating progenitors of the mouse fetal intestinal epithelium. Cell Rep 5, 421–432. 10.1016/j.celrep.2013.09.005

Nakayama, M., Oshima, M., 2019. Mutant p53 in colon cancer. Journal of Molecular Cell Biology 11, 267–276. 10.1093/jmcb/mjy075

Neph, S., Kuehn, M.S., Reynolds, A.P., Haugen, E., Thurman, R.E., Johnson, A.K., Rynes, E., Maurano, M.T., Vierstra, J., Thomas, S., Sandstrom, R., Humbert, R., Stamatoyannopoulos, J.A., 2012. BEDOPS: high-performance genomic feature operations. Bioinformatics 28, 1919–1920. 10.1093/bioinformatics/bts277

Nieto, M.A., 2013. Epithelial Plasticity: A Common Theme in Embryonic and Cancer Cells. Science 342. 10.1126/science.1234850

Nieto, M.A., Sargent, M.G., Wilkinson, D.G., Cooke, J., 1994. Control of cell behavior during vertebrate development by Slug, a zinc finger gene. Science 264, 835–839. 10.1126/science.7513443

Ocaña, O.H., Córcoles, R., Fabra, Á., Moreno-Bueno, G., Acloque, H., Vega, S., Barrallo-Gimeno, A., Cano, A., Nieto, M.A., 2012. Metastatic Colonization Requires the Repression of the Epithelial-Mesenchymal Transition Inducer Prrx1. Cancer Cell 22, 709–724. 10.1016/j.ccr.2012.10.012

Oliemuller, E., Newman, R., Tsang, S.M., Foo, S., Muirhead, G., Noor, F., Haider, S., Aurrekoetxea-Rodríguez, I., Vivanco, M. dM, Howard, B.A., 2020. SOX11 promotes epithelial/mesenchymal hybrid state and alters tropism of invasive breast cancer cells. eLife 9, e58374. 10.7554/eLife.58374

Parikh, K., Antanaviciute, A., Fawkner-Corbett, D., Jagielowicz, M., Aulicino, A., Lagerholm, C., Davis, S., Kinchen, J., Chen, H.H., Alham, N.K., Ashley, N., Johnson, E., Hublitz, P., Bao, L., Lukomska, J., Andev, R.S., Björklund, E., Kessler, B.M., Fischer, R., Goldin, R., Koohy, H., Simmons, A., 2019. Colonic epithelial cell diversity in health and inflammatory bowel disease. Nature 567, 49–55. 10.1038/s41586-019-0992-y

Phipps, A.I., Buchanan, D.D., Makar, K.W., Win, A.K., Baron, J.A., Lindor, N.M., Potter, J.D., Newcomb, P.A., 2013. KRAS-mutation status in relation to colorectal cancer survival: the joint impact of correlated tumour markers. Br J Cancer 108, 1757–1764. 10.1038/bjc.2013.118

Pobbati, A.V., Kumar, R., Rubin, B.P., Hong, W., 2023. Therapeutic targeting of TEAD transcription factors in cancer. Trends in Biochemical Sciences 48, 450–462. 10.1016/j.tibs.2022.12.005

Robinson, J.T., Thorvaldsdóttir, H., Winckler, W., Guttman, M., Lander, E.S., Getz, G., Mesirov, J.P., 2011. Integrative Genomics Viewer. Nat Biotechnol 29, 24–26. 10.1038/nbt.1754

Robinson, M.D., McCarthy, D.J., Smyth, G.K., 2010. edgeR: a Bioconductor package for differential expression analysis of digital gene expression data. Bioinformatics 26, 139–140. 10.1093/bioinformatics/btp616

Roper, J., Tammela, T., Akkad, A., Almeqdadi, M., Santos, S.B., Jacks, T., Yilmaz, Ö.H., 2018. Colonoscopy-based colorectal cancer modeling in mice with CRISPR-Cas9 genome editing and organoid transplantation. Nat Protoc 13, 217–234. 10.1038/nprot.2017.136

Sato, T., Clevers, H., 2013. Growing self-organizing mini-guts from a single intestinal stem cell: mechanism and applications. Science 340, 1190– 1194. 10.1126/science.1234852

Shannon, P., Markiel, A., Ozier, O., Baliga, N.S., Wang, J.T., Ramage, D., Amin, N., Schwikowski, B., Ideker, T., 2003. Cytoscape: A Software Environment for Integrated Models of Biomolecular Interaction Networks. Genome Res. 13, 2498–2504.10.1101/gr.1239303

Siegel, R.L., Giaquinto, A.N., Jemal, A., 2024. Cancer statistics, 2024. CA: A Cancer Journal for Clinicians 74, 12–49. 10.3322/caac.21820

Stuart, T., Butler, A., Hoffman, P., Hafemeister, C., Papalexi, E., Mauck, W.M., Hao, Y., Stoeckius, M., Smibert, P., Satija, R., 2019. Comprehensive Integration of Single-Cell Data. Cell 177, 1888-1902.e21. 10.1016/j.cell.2019.05.031

Szklarczyk, D., Kirsch, R., Koutrouli, M., Nastou, K., Mehryary, F., Hachilif, R., Gable, A.L., Fang, T., Doncheva, N.T., Pyysalo, S., Bork, P., Jensen, L.J., von Mering, C., 2023. The STRING database in 2023: protein–protein association networks and functional enrichment analyses for any sequenced genome of interest. Nucleic Acids Research 51, D638–D646. 10.1093/nar/gkac1000

Tarasov, A., Vilella, A.J., Cuppen, E., Nijman, I.J., Prins, P., 2015. Sambamba: fast processing of NGS alignment formats. Bioinformatics 31, 2032– 2034. 10.1093/bioinformatics/btv098

Thiery, J.P., Acloque, H., Huang, R.Y.J., Nieto, M.A., 2009. Epithelial-Mesenchymal Transitions in Development and Disease. Cell 139, 871–890. 10.1016/j.cell.2009.11.007

Tiberi, S., Crowell, H.L., Samartsidis, P., Weber, L.M., Robinson, M.D., 2023. distinct: A novel approach to differential distribution analyses. The Annals of Applied Statistics 17, 1681–1700. 10.1214/22-AOAS1689

Virchow, R.L.K., 1859. Die Cellularpathologie in ihrer Begründung auf physiologische und pathologische Gewebelehre. A. Hirshcwald.

Vuik, F.E., Nieuwenburg, S.A., Bardou, M., Lansdorp-Vogelaar, I., Dinis-Ribeiro, M., Bento, M.J., Zadnik, V., Pellisé, M., Esteban, L., Kaminski, M.F., Suchanek, S., Ngo, O., Májek, O., Leja, M., Kuipers, E.J., Spaander, M.C., 2019. Increasing incidence of colorectal cancer in young adults in Europe over the last 25 years. Gut 68, 1820–1826. 10.1136/gutjnl-2018-317592

Weinberg, R.A., 1996. How Cancer Arises. Scientific American 275, 62–70.

Wickham, H., 2016. ggplot2, Use R! Springer International Publishing, Cham. 10.1007/978-3-319-24277-4

Yang, C., Croteau, S., Hardy, P., 2021. Histone deacetylase (HDAC) 9: versatile biological functions and emerging roles in human cancer. Cell Oncol. 44, 997–1017. 10.1007/s13402-021-00626-9

Youssef, K.K., Narwade, N., Arcas, A., Marquez-Galera, A., Jiménez-Castaño, R., Lopez-Blau, C., Fazilaty, H., García-Gutierrez, D., Cano, A., Galcerán, J., Moreno-Bueno, G., Lopez-Atalaya, J.P., Nieto, M.A., 2024. Two distinct epithelial-to-mesenchymal transition programs control invasion and inflammation in segregated tumor cell populations. Nat Cancer 1–21. 10.1038/s43018-024-00839-5

Yu, G., Wang, L.-G., Han, Y., He, Q.-Y., 2012. clusterProfiler: an R package for comparing biological themes among gene clusters. OMICS 16, 284–287. 10.1089/omi.2011.0118

Yu, G., Wang, L.-G., He, Q.-Y., 2015. ChIPseeker: an R/Bioconductor package for ChIP peak annotation, comparison and visualization. Bioinformatics 31, 2382–2383. 10.1093/bioinformatics/btv145

Zhang, W.C., Shyh-Chang, N., Yang, H., Rai, A., Umashankar, S., Ma, S., Soh, B.S., Sun, L.L., Tai, B.C., Nga, M.E., Bhakoo, K.K., Jayapal, S.R., Nichane, M., Yu, Q., Ahmed, D.A., Tan, C., Sing, W.P., Tam, J., Thirugananam, A., Noghabi, M.S., Huei Pang, Y., Ang, H.S., Mitchell, W., Robson, P., Kaldis, P., Soo, R.A., Swarup, S., Lim, E.H., Lim, B., 2012. Glycine Decarboxylase Activity Drives Non-Small Cell Lung Cancer Tumor-Initiating Cells and Tumorigenesis. Cell 148, 259–272. 10.1016/j.cell.2011.11.050

Zhao, L., Song, W., Chen, Y.-G., 2022. Mesenchymal-epithelial interaction regulates gastrointestinal tract development in mouse embryos. Cell Rep 40, 111053. 10.1016/j.celrep.2022.111053

