## Supplementary tables for "An oncoembryology approach uncovers SoxC-driven regulation of colon development and cancer"

**Table S1.**
**Primers used for genotyping SoxC fl/fl and R26-CreERT2 mouse alleles.**

| | | |
|---|---|---|
| Sox4 F | GAAGGAGGCGGAGAGTAGACGG | fl 520 |
| Sox4 R | CTAAGCTCAACACAAATGCCAAACGC | WT 419 |
| Sox11 F | TTCGTGATTGCAACAAAGGCAGAG | fl ca. 400 |
| Sox11 R | GCTCCCTGCAGTTTAAGAAATCGG | WT 305 |
| Sox12 F | GGAGAACAGATGGGCAGCG | fl 516 |
| Sox12 R | GGCCAGTAGAGCTCCTCCG | WT 468 |
| R26-CreERT2 1 | AAAGTCGCTCTGAGTTGTTAT | Mut 825<br>WT 603 |
| R26-CreERT2 2 | GGAGCGGGAGAAATGGATATG | |
| R26-CreERT2 3 | CCTGATCCTGGCAATTTCG | |

**Table S2.**
**Primary Antibodies used**

| Protein | Catalogue nr. | Dilution |
|---|---|---|
| Trop2 (*Tactsd2*) | Invitrogen, PA5-47074 | 1:100 |
| Marckls1 | Atlas Antibodies HPA030528 | 1:100 |
| Sox11 | Atlas Antibodies HPA000536 | 1:100 IF, 1:500 ChIP |
| Sox4 | Diagenode C15310129 | 1:100 ChIP |
| Epcam | Abcam ab213500 | 1:200 |
| Klf4 | Biotechne AF3158 | 1:200 |
| Tead2 | Atlas Antibodies HPA066292 | 1:50 |
| Mdk | Atlas Antibodies HPA057126 | 1:100 |
| E-cadherin | Biotechne AF748 | 1:200 |
| Ki67 | Abcam ab15580 | 1:1000 |
| Klf5 | R&D Systems, AF3758 | 1:100 |

**Table S3.**
**List of qPCR primers**

| | |
|---|---|
| bActin F | GATCTGGCACCACACCTTCT |
| bActin R | GGGGTGTTGAAGGTCTCAAA |
| *Klf4* F | CTCTGCTCCCGTCCTTCTC |
| *Klf4* R | TTGAACTCCTCGGTCTCCCT |
| *Klf5* F | TCTGGAGAAGCGACGTATCC |
| *Klf5* R | CTCCCAGGTGCACTTGTAGG |
| *Mdk* F | GAAGAAGGCGCGGTACAATG |
| *Mdk* R | CTGGCCTCCTGACTTAGTCC |
| *Tead2* F | TGCCTTCTTCCTCGTCAAGT |
| *Tead2* R | AGGTGAGTGTCATGAGCTCC |
| *Sox4* F | GATGCGTTTGGCATTTGTGT |
| *Sox4* R | TCTCCAGCTGCAAGGACAAG |
| *Sox11* F | GACGACCTCATGTTCGACCT |
| *Sox11* R | GGGGAACTCGAAGTGGGAA |
| *Sox12* F | GGCTCCTCCCTAAGTCCATC |
| *Sox12* R | GGTCTCTCCTCAAGCCAAGT |

**Table S4.**
**List of primers used for cloning purposes.**

| | |
|---|---|
| Tead2 enh. F | TGCACTCGAGACCCACACTTCTATACACACACAAGTA |
| Tead2 enh. R | TCGAAAGCTTCAAGACAGACGGACATCTCTAATCT |
| Mdk silencer F | GCTAGCCCGGGCTCGAGTGCCACATCAGAGACCCTTC |
| Mdk silencer R | CCATTATATACCCTCTAGAGTCTAGATCTCGAGGTGCTGCATCCTAGGGGTAA |
| Sox11-CDS F | TCGAGTCGACTCGACATATGACTCTAGAGGATCCATGGTGCAG |
| Sox11-CDS R | GCTGATCAGCTTCTGCTCATCCATATACGTGAACACCAGGT |
| Myc-P2A-GFP F | GAGCAGAAGCTGATCAGCGAGGAGGACCTGGGAAGCGGAGCTACTAACTTCA |
| Myc-P2A-GFP R | CATATGTCGAATTTAAATTCGAGTCGACTTACACCTTCCTCTTCTTCTTAG |
| pLenti-Lifeact-EGFP F | TCGAATTTAAATTCGACATATGAATCAACCTCTGGATTACAAAATTTGTGAAAGATT |
| pLenti-Lifeact-EGFP R | CATATGTCGAGTCGACTCGAAGGTCAAAACAGCGTGGATGG |
| Sox4-CDS-3myc-P2A F | AGGTCGACTCTAGAGGATCGAATTCCAAGCCGGGGCCATGGTA |
| Sox4-CDS-3myc-P2A R | TCAGGTCGGCCACGCCCATGAATTCAGGTCCAGGGTTCTCCTCC |
| Klf4-CDS F | CGCTGTTTTGACCTCCATAGAAGATTCTAGAGCCACCATGAGGCAGCCACCTGG |
| Klf4-CDS R | TCCGATTTAAATTCGAATTCGCTAGCTCTAGATTAAAAGTGCCTCTTCATGTGTAAGGC |
| Tead2 enh. Mutation F | GAAAGTTGCCGGGAGTGGGAAAAGTGGGAG |
| Tead2 enh. Mutation R | CTCCCACTTTTCCCACTCCCGGCAACTTTC |
